# Modeling EEG Resting-State Brain Dynamics: A proof of concept for clinical studies

**DOI:** 10.1101/2023.04.15.536991

**Authors:** Sonja A. Kotz, Aland Astudillo, David Araya, Simone Dalla Bella, Nelson Trujillo-Barreto, Wael El-Deredy

## Abstract

Functional brain imaging has shown that the awake brain, independent of a task, spontaneously switches between a small set of functional networks. How useful this dynamical view of brain activity is for clinical studies, e.g., as early markers of subsequent structural and/or functional change or for assessing successful training or intervention effects, remains unclear. Core to addressing this question is to assess the robustness and reproducibility of the analysis methods that model, characterize, or infer the features of brain dynamics, and the accuracy by which these features represent and classify specific cognitive or altered cognitive states. This is particularly key given inter- and intra-individual variability and measurement noise. Here we used resting-state EEG from persons with Parkinson’s Disease (PD) and healthy matched controls to systematically assess the reliability, robustness, and sensitivity of Hidden semi-Markov models (HsMM). These models are an example of model-based probabilistic methods for Brain-State allocations that are estimated from observed data. The method estimates model parameters, if the M/EEG recording or observations, over the scale of minutes, are emissions from hidden states that persist over short durations, before switching or transitioning to other states. We introduce an analysis pipeline that leads to sets of reproducible features of neurophysiological dynamics at the individual level. These features can be used as discriminatory variables to classify individuals and to evaluate the effect of non-pharmacological training schemes like in the current example a music-gait exercise program for Parkinson’s Disease. Given the method stochasticity and the data variability, we emphasize the importance of repeating the analysis to reliably identify brain states and their dynamical trajectories that subsequently can be related to individualized variables.

## 1. Introduction

Early reorganization of functional networks in neurodegeneration should manifest as changes in ongoing brain dynamics, even in the absence of symptoms (Babiloni et al., 2013; Hatz et al., 2015; Tsvetanov et al., 2021). Functional brain networks reorganize, either by slow changes in the relative contribution or rewiring of brain areas (plasticity) or via fast modulation of their causal interactions (effective connectivity). This reorganization compensates for early neurodegenerative changes, and thus can mask or delay observable deficits (Chen et al., 2017). Thus, monitoring brain function as a series of static task-specific states may not be the most effective way to denote early markers for neurodegeneration. Rather, important clues may lay in the path or trajectory by which functional brain networks evolve and transit over time, even in the absence of a task, and the time it dwells in each functional network (Hansen et al., 2015). Indeed, functional imaging studies have shown that the awake brain spontaneously switches between a small set of functional networks (Vidaurre et al., 2017). Yet, estimating the evolution of the non-stationary and non-linear brain is methodologically challenging. There is a growing interest in the development of methods that explicitly model the brain’s dynamical regimes, across multiple temporal length-scales, over which the brain networks are assumed to switch, from a few milliseconds to seconds (Baker et al., 2014). These methods aim to identify and characterize transiently stable and recurrent activity patterns in the ongoing M/EEG (magneto/electroencephalography) at topographical (Michel and Koenig, 2018; Becker et al., 2020; Bolton et al., 2020), sources (Baker et al., 2014; Woolrich et al., 2013; Trujillo-Barreto et al., 2019), or functional connectivity levels (Olier et al., 2013; Vidaurre et al., 2018; Kottaram et al., 2019). However, it remains unclear whether the features of ongoing brain dynamics that these methods extract can be clinically relevant and sufficiently sensitive in detecting abnormalities or monitoring disease progression.

An example of such methods is a class of probabilistic models where the measured M/EEG data are generated from unobservable hidden brain states that persist over time before instantaneously switching to other states (Fig. 1). Here we provide evidence that features of the dynamics and the methodology to infer them would be sufficiently reliable and robust to be applied in clinical studies (Khanna et al., 2015; O’Neill et al., 2018; Preti et al., 2017). For example, we assume that early stages of neurodegeneration can be detected and differentiated as alterations in the dynamical repertoires that the brain explores, well before symptoms are reported (Wang et al., 2021; Zhao et al., 2022). These alterations would be reflected either as the probability of the brain switching between states, the duration it spends in a particular state, or in terms of the sequence or path it takes between states. This is consistent with the observation that the ageing human brain undergoes dynamical changes to accommodate sensory fragility (Cabeza et al., 2018). Therefore, transitions between the hidden brain states and their durations (dwell times) may reflect properties of the brain that could be used as markers of disease and disease progression (Van de Ville et al., 2010). The hidden states could be regarded as abstract operation modes or as network configurations, each of which generates the observed data. However, the physiological or biophysical interpretation of those hidden states remains unclear (Kringelbach and Deco, 2020).

**Fig. 1.**
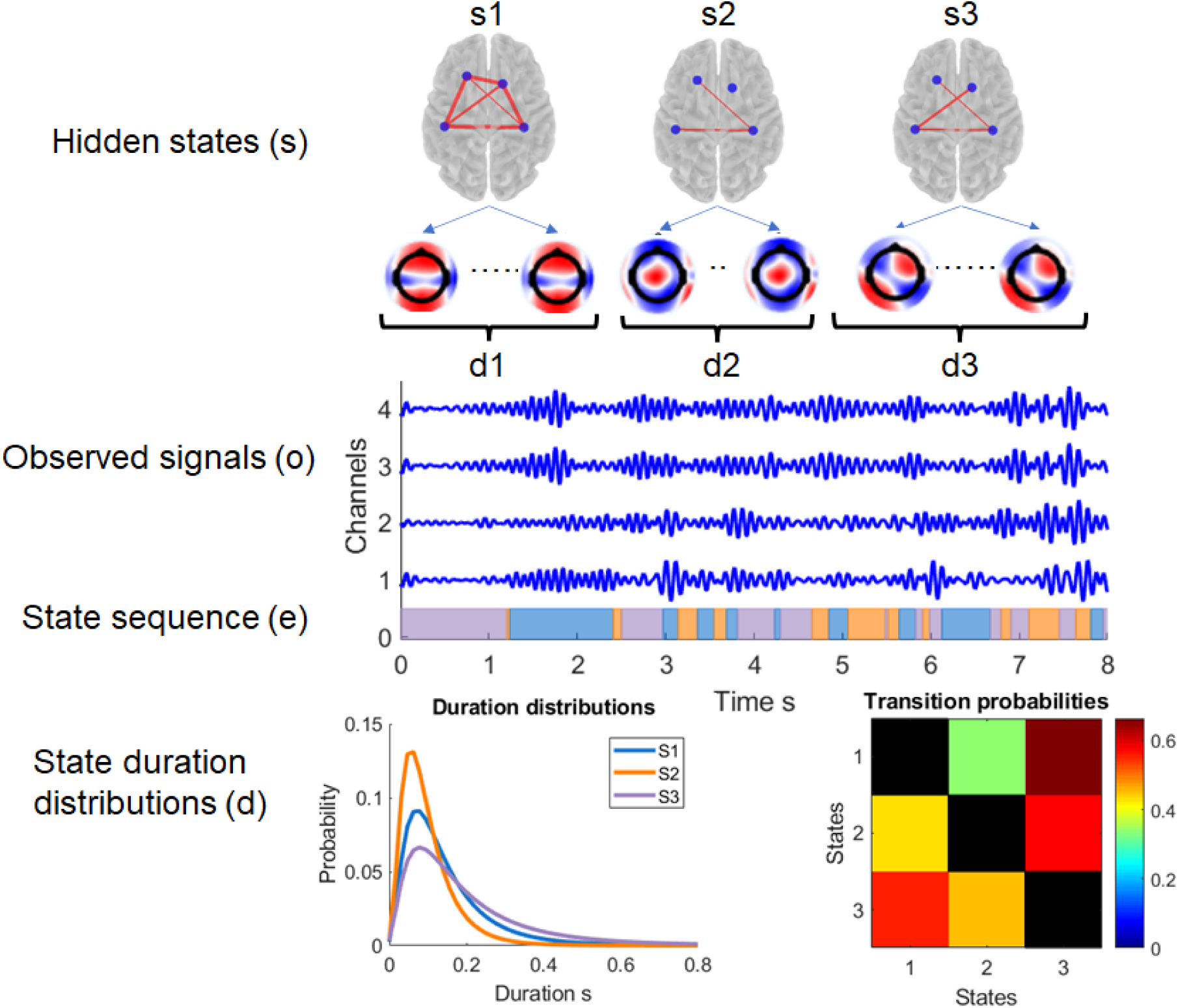
Conceptual representation of modeling dynamic brain state allocation using hidden semi-Markov models (HsMM). Observed (recorded) signals at the level of the EEG channels or source activity are assumed to be generated as continuous observations/emissions (**o**) from hidden discrete states (**s**) corresponding to different functional network configurations whose activation persists in time (**d**), before transitioning probabilistically to other states according to transition matrix (**T**). HsMM is a probabilistic model of how the data was generated, according to this process. Modeling entails using the observed data to recover or estimate the generating parameters, i.e., the characteristics of how a particular sequence of data has been generated. These are the distributions of the emissions and durations, and the transition probability matrix. In doing so, the model classifies the observed time series in according to the sequence of states that generated it (**e**).

Here we focus on the reliability of the analysis methods in extracting credible information dynamics, particularly given the intra- and inter-subject variability in clinical data and beyond. We aim to show that the applied methods are reproducible and robust in obtaining similar results from repeated data modeling. This is in line with the current challenges to make the application of models useful in neuroscience and health care, where reproducible and replicable results are required. This includes ensuring ranges of result reproducibility and modeling given the uncertainty of the data and the stochastic nature of the analysis methods (Milkowski et al., 2018; Beam et al., 2020). Uncertainty about the methods stems from their sensitivity to the initial conditions that are usually chosen at random and not guaranteed to converge to a globally optimal set of dynamical features.

As a proof of concept for the clinical utility of modeling brain dynamics, we used resting state EEG (RS-EEG) data from persons with Parkinson’s Disease (PD) and their matched healthy controls. Emerging evidence suggests that PD is associated with structural and functional brain network alterations (Fornito et al., 2015) due to progressive dopamine depletion (Roheger et al., 2018) that subsequently drive observable EEG dynamics, different from fluctuations observable in the healthy aging brain (Hohlefeld et al., 2015). EEG spectral analysis shows alteration in alpha power in PD patients (Caviness et al., 2016) and correlates with clinical symptoms (Aarsland et al., 2009; Babiloni et al., 2011; Hassan et al., 2017; Yılmaz et al, 2020). Additionally, a microstate analysis study on PD levodopa effects showed that task-free EEG patterns changed in specific microstates profiles and durations (Serrano et al., 2018). Taken together, PD presents different brain structural and functional brain networks alterations that may relate to progressive motor and cognitive dysfunctions. We hypothesized that brain dynamics are also different in persons with PD. This difference should show in spontaneous EEG as changes in the dynamical features estimated by a brain-state allocation tool; if this tool is sufficiently reliable, the dynamical features should be sensitive to the neural dynamic changes induced by training or intervention activities.

## 2. Materials and methods

### 2.1 Participants and data acquisition

We analyzed RS-EEG collected next to behavioral data (for report on the behavioral data see Bella et al., 2017). The study included 14 Idiopathic PD patients and 14 matched healthy controls. Persons with PD participated in an auditory-motor synchronization exercise, comprising musically cued movement. This procedure has shown to change behavioral and motor performance in PD patients (Bella et al., 2017). The study was approved by the Ethics Committee of the University of Leipzig, Leipzig, Germany. Spontaneous ongoing (resting-state) EEG (RS-EEG) activity was recorded before and after the exercise program – thereafter referred to as *PD-pre-* and *PD-post exercise program*. 2-minute blocks of ongoing RS-EEG data were recorded while alternating between eyes open and eyes closed conditions. EEG was recorded from 60 10-20 scalp electrodes, referenced to the left mastoids (Brain Amps, Brain Vision, NC, USA), at 500 Hz, with impedances kept below 20 kΩ. Eye movements were recorded with 4 additional electrodes positioned around the eyes.

### 2.2 Data pre-processing and feature extraction for modeling

Two minutes of eyes closed RS-EEG data were used as eyes closed data have a better signal-to-noise ratio for analyzing activity in the alpha band. The data were pre-processed using the EEGLab toolbox (Delorme and Makeig, 2004), running in MATLAB (MathWorks Inc., Natick MA, United States). The data cleaning was performed independently for each participant. Raw RS-EEG signals were filtered between 0.1 Hz -120 Hz and detrended. Ocular, cardiac, motion, and main interference artifacts were corrected using an independent component analysis (ICA) as in (Baker et al., 2014; Hunyadi et al., 2019). The cleaned data were referenced to the average of the scalp electrodes.

We followed the modeling pipeline of (Baker et al., 2014) (see Fig. 2). The cleaned RS-EEG data were filtered to the alpha band (7 - 14 Hz). The amplitude envelope of the oscillatory activity at each channel was then computed using the Hilbert transform. Each channel was demeaned and normalized by the variance over channels to guarantee the same scale for the envelope data of all participants. The standardized envelope time series from all participants were concatenated, forming an ensemble of 60 channels by a time series of length 14 x 3 participants (14 control participants and 2 x 14 for the PD patients pre- and post-exercise program) x 120 s x 500 samples/s. The dimensionality of the concatenated ensemble was reduced using Principal Component Analysis (PCA) keeping 20 principal components (PCs) explaining 86% of the variance. The Hilbert envelope-PCA transformed data were the input to the HsMM modeling algorithm.

**Fig. 2.**
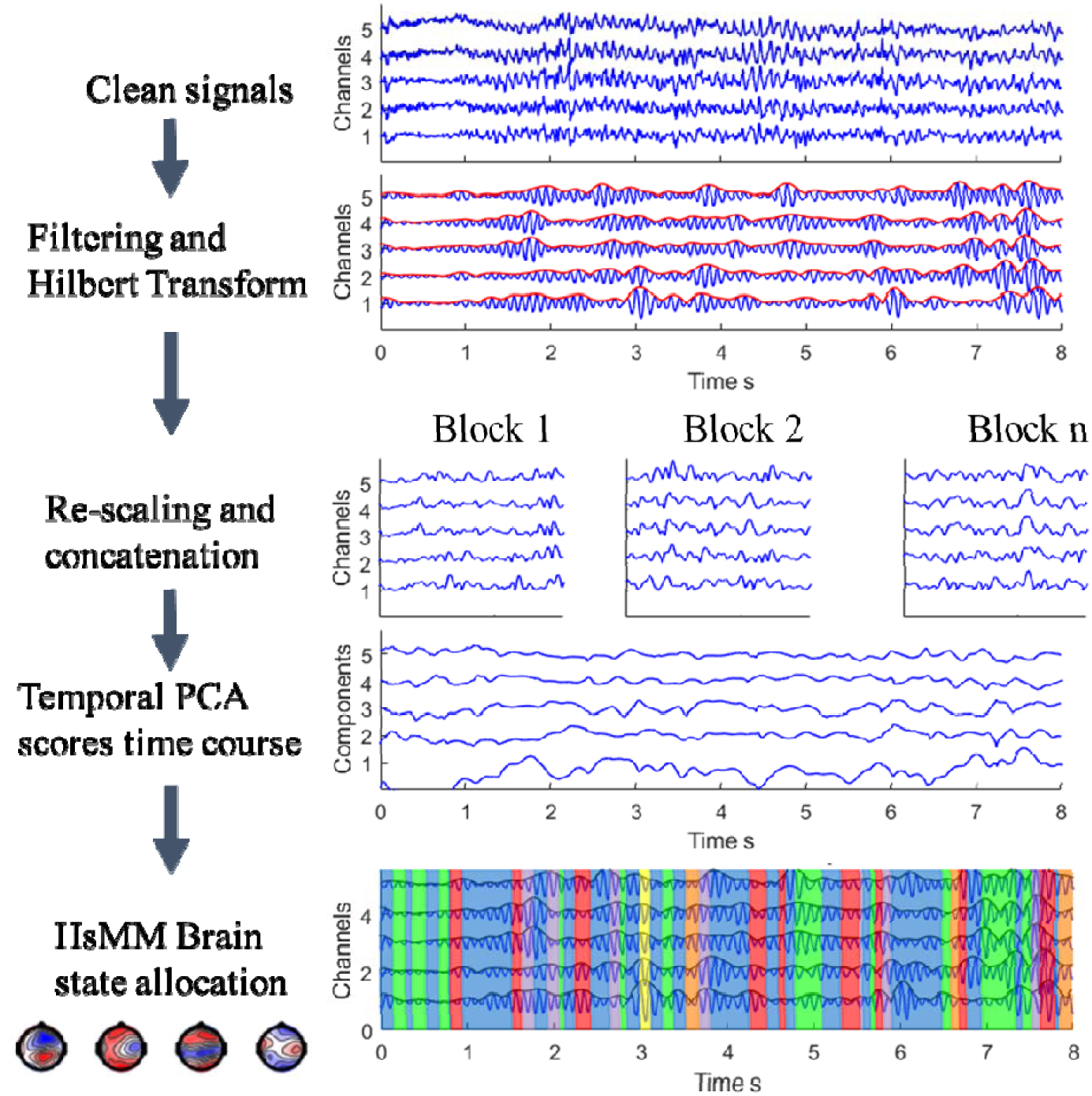
Feature extraction and modeling pipeline. Artefact free RS-EEG data are band filtered (here we focused on the alpha frequency band). The amplitude envelope of the band of interest is extracted using the Hilbert Transform. Data from individual recording blocks of participants are normalized, scaled, and then concatenated. Data reduction is achieved by taking the top principal components and by down sampling. HsMM is applied to estimate the state topographies, state sequence, and transition matrix. Using the state sequence, empirical metrics are obtained: fractional occupancies (FO) and dwell times (DT).

### 2.3 Dynamical state allocation with Hidden semi-Markov Modeling

A Hidden Semi-Markov Model (HsMM) is an extension of the classic Hidden Markov Model (HMM) where state durations are explicitly defined and estimated (Fig. 1). The need for an explicitly specified state duration model follows from the observation that the brain probably obeys a log-normal distribution (Buzsaki and Mizuseki, 2014). This permits the estimation of state-durations consistent with what is observed empirically. Here, we followed the HsMM implementation of Trujillo-Barreto et al. (2019). This implies that in each state, instead of transitioning at every time-point as in the classical HMM, the emission model generates a data sequence whose length is determined by the duration distribution of that state, before transitioning to a new state. Self-transitioning is not allowed. In essence, HMM/HsMM are detectors of change points in the statistical properties of the M/EEG time series. In other words, the model segments the time series into chunks with shared spatio-temporal statistical properties. In the case of the current HsMM, these data chunks have state durations that follow a log-normal distribution.

In HMM/HsMM the observed/measured data is modeled as continuous emissions from discrete hidden, unobservable brain states (Fig. 1). The states are abstract but could be interpreted as stable configuration of functional networks, evolving over time. The modeling/learning algorithm estimates the probability of the hidden states, the transition probabilities between them, and the distribution of emissions (the observations) from each state (here assumed to be multivariate normal, MVN distribution). HsMM circumvents the limitation of HMM that implies (incorrectly) that the duration the brain spends in each state follows a geometric distribution, by explicitly defining a more appropriate duration model, here assumed to be a log-normal (Trujillo-Barreto et al., 2019). However, there is no consensus on the correct number of states to be modeled (Quinn et al., 2018; Vidaurre et al., 2017). Here, we used a Bayesian model comparison to identify an optimal number, in repeated runs.

### 2.4 State sequence and state activity maps

#### State sequence

Refers to a labeling or designation of each point in the time series to the most probable hidden state. After the model is trained, the data points, or new data, are decoded to obtain the probability of each state being responsible for generating each observation. The time points are allocated to the state with maximum probability of emission given that state.

#### State topographies

As mentioned in Section 2.2, the training input to the HsMM, i.e., the observations, is the PCA dimensionality reduced envelope concatenated data. Thus, the estimated emissions from the HsMM remain in the lower dimension PC space where each state has a MVN distribution associated with a mean vector. To project back to scalp maps associated with each state, we used the state mean and the PCA coefficients and we projected back onto the sensor space. The resulting coefficients corresponded to maps of state activity patterns. This approach is equivalent to the general linear model (GLM) used by Baker et al., (2014).

To illustrate the link between the state topography at the electrode level and the underlying networks, we estimated the neural sources of the state topographies using ELORETA toolbox (exact low-resolution electrical tomography analysis; Pascual-Marqui, 2007). These are shown in the *Supplementary Material*.

### 2.5 Extracting individual metrics

Like (Baker et al., 2014), we used state sequence derived metrics to characterize results on an individual basis. Three metrics were obtained from the state sequences: the fractional occupancy (FO), the mean dwell time (DT) per state, and the probability of transition between states. The FO is defined as the fraction of time that the observations remain in each state relative to the entire duration of the recorded data. The DT of a state is the average time spent in a state where the average is taken over the number times the state has been visited.

We calculated the *FO, DT*, and *transitions* for all participants in each group and compared their values between groups. These metrics were obtained empirically from the state sequences. In the case of the DT, its distribution was fitted using a log-normal for each participant/block, obtaining 2 parameters: mean and variance.

### 2.6 Modeling experiments

#### Exp. 1. Choosing the number of states and individual participants

The aim of this experiment was to demonstrate the convergence of a consistent number of states over repeated runs. Data from one participant were used to train 50 models, while changing the number of states from 4 to 10, with a Bayesian model selection using the Free-Energy (FE) (Trujillo-Barreto et al., 2019), and determining the most probable number of states. This experiment was repeated over ten randomly chosen participants to estimate the number of states that was then applied to subsequent group experiments (Supp. Fig. 1).

#### Exp. 2. Reproducibility of state allocation over repeated runs

This experiment should demonstrate the robustness of HsMM modeling over repeated runs. This was done at the single participant and the group level. HsMM model instantiations were trained on data from a single participant. Twenty independent model instantiations were established where each model had a different random state sequence initialization. Consistency among model instantiations was assessed by testing the resulting model parameters and the distribution over state sequence metrics. FO and state maps were obtained. State maps for model instantiations were compared.

Concatenated transformed data from 14 healthy participants and 14 PD pre-music-gait exercise program was used for training 20 independent models, for a fixed number of ***K*** = 9 states. Model parameters, state sequences, state maps, and state sequence metrics were obtained for each model instantiation. Distributions of state sequence metrics were compared across model instantiations.

Consistency at the individual level as well as group level metrics were assessed by comparing the variability between resulting values (Cohen, 2014). Results for each comparison show the variability across different model instantiations. We assessed differences between state topographies. In a similar way, we compared FOs and DTs across participants and groups.

The reproducibility of the state maps was evaluated based on the correlations between them over instantiations. The reliability of the models was expressed in terms of distributions of the DTs of all models. The aim was to test if each characterization was robust and reproducible in terms of pipeline steps and configuration settings.

#### Exp. 3. Classification of individuals based on the dynamical features

This experiment tested if the individual dynamical features, FO, DT, estimated from the HsMM model, carry sufficient information to characterize and differentiate healthy participants and PD patients. Additionally, We compared HsMM features against a classifier based on the standard power spectral analysis in the alpha band (Chu et al., 2021; Mei et al., 2021). We measured the consistency of HsMM outputs as features to characterize individuals and groups. We built a simple log-regression GLM classifier (Yang and Huang, 2022), on either FO, DT, or their combination from each data block of the control participants and PD patients’ pre-music-gait exercise program. We used cross validation leave one out to test the classification performance. The probability of each data block being classified as healthy controls (class label = 0), or PD patients (class label = 1) was compared across data groups and model instantiations. Estimation of the Receiver Operator Characteristic (ROC) curve, the area under the curve (AUC), and comparison to a classifier based on power spectral alpha band values are reported in the *Supplementary Material*.

#### Exp. 4. Individual difference in response to music-cued movement exercise program

This experiment tested whether the individual dynamical features of the PD patients, estimated from the HsMM model, carried sufficient information to track changes in the individual following the music-cued movement exercise program.

EEG data from the PD patients’ post-music-cued exercise were not used to train the HsMM model. Instead, data were decoded by the already trained HsMM model on the healthy controls and the PD patients (pre-exercise program) data. The decoding procedure is a type of inference, where the trained model extracts the dynamical features, FO, DT, and transitions of the previously unseen data. We assumed that the model of healthy controls plus the PD patients (pre-exercise program) captures the range of dynamics of the sample, and that the PD data post-music-cued exercise program would represent variation on these dynamics, where some patients would look more comparable to healthy controls, some would remain the same, while others might not show any improvement after the training program.

The PD patients post-exercise program data was transformed using the described approach in previous section. The PCA envelopes of PD patient post-exercise program data was decoded used the pretrained model. The decoded state sequences from the PD patients post-exercise program were used to extract state metrics: FO and DT, which were then used as inputs to the classifier in Experiment 3, to re-classify the PD patients post-exercise program and see the effects of the intervention in comparison to the PD patients’ pre-exercise program.

### 2.7 Statistical analyses

Statistical analyses were carried out in MATLAB (MathWorks Inc., Natick MA, United States), JAMOVI (The Jamovi Project, 2022, Sydney, Australia) and SPSS for Windows, version 23.0 (IBM Inc., Chicago, IL, United States).

#### Variability of state maps across model runs/instantiations

The variability of state maps across model instantiations at the individual and the group level was tested by measuring the distance between state maps within and between states. We compared the average difference between common states versus different states using t-tests. A statistically significant value was set to p < 0.05.

#### Differences in FO and DT across model runs/instantiations, and between groups

Differences in FO and DT were compared across model instantiations within and between individuals of the healthy control participants and persons with PD. The difference of the average FO and the average DT between each group was tested using t-tests. For the group comparison in one model, the different FO and DT averages were verified by a t-test for repeated measures with bootstrapping (***n*** = 1000).

#### Testing classification scores over across model runs/instantiations

The output of the classifier is a score that measures how likely the input values of FO and DT belong to the PD or the healthy control group. Variability of the classification scores was assessed among independent model instantiations. The score values were compared across models for each participant by means of an analysis of variance. The same evaluation was performed for the classification scores of PD post-exercise data. We used the Wilcoxon test to test the variance.

## 3. Results

### Exp. 1. Choice of number of states, individual participants

The model implemented by Trujillo-Barreto and colleagues (2019) uses the variational Free-Energy (FE) as the cost-function to estimate the model. We used FE to compare between models. Fig. 3 shows the convergence of FE over repeated runs in one participant. The figure highlights the variability in convergence due to different initial conditions, and the importance of repeated modeling to arrive at a reliable conclusion about the appropriate number of states. This is important for short data sequences or small sample sizes. Due to the number of states and parameters, this model approach needs enough data points to characterize each participant’s data.

**Fig. 3.**
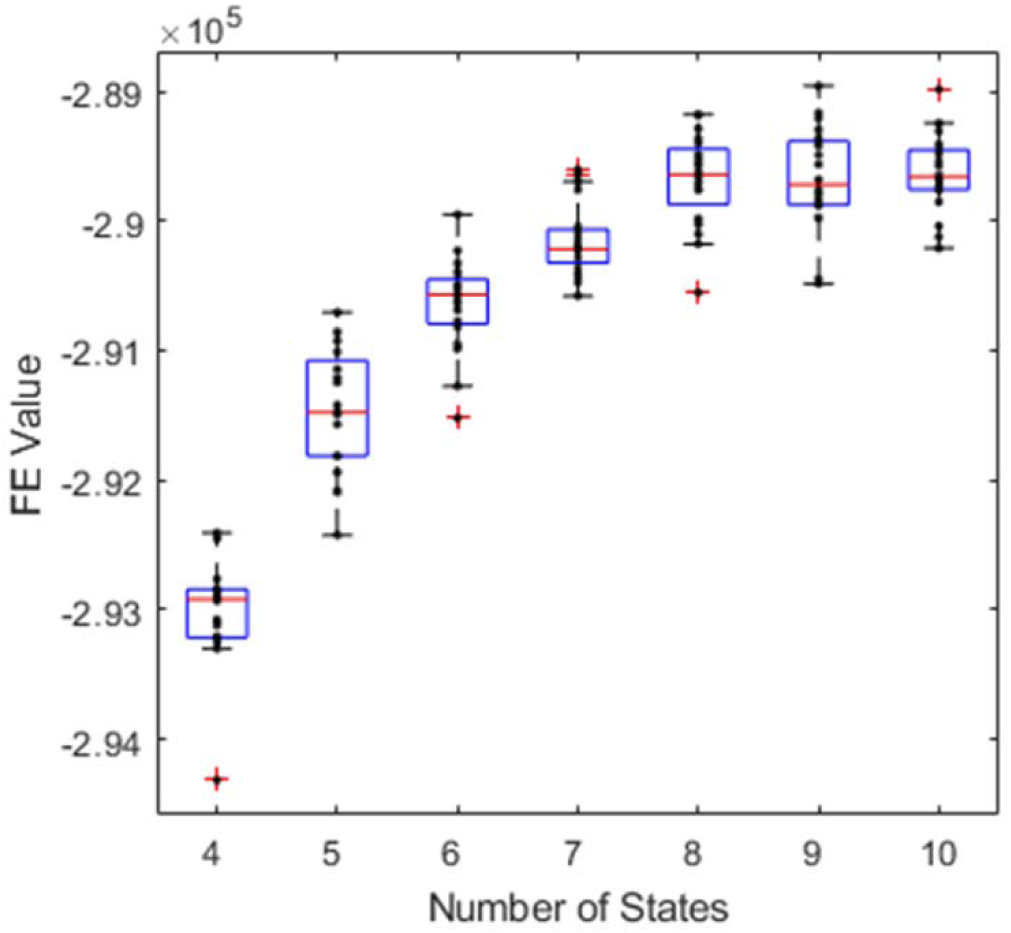
Reproducibility of convergence to the number of states. Twenty repetitions modeling 2 min of RS-EEG data from a single PD patient pre-exercise program. The Free-Energy (FE) was used as model comparison criterion to choose the most likely number of states. Repeated runs result in different FE paths depending on the (random) initial conditions. The figure highlights the need for repeated runs from different initial conditions to arrive at a robust conclusion about the number of states, particularly when the sequence of modeled EEG data is short, as in the case here (2 min). The figure shows a majority convergence to ***K*** = 9 states (**37** times out of 50).

Additionally, this experiment was repeated over ten randomly chosen participants to estimate the optimal number of states that was then applied to subsequent group experiments (Supp. Fig. 1). The most common outcome was ***K*** = 9 states, which was the fixed value used for the subsequent group level analysis.

### Exp. 2. Reproducibility of state allocation over repeated runs

First, we evaluated the reproducibility at the single participant level. Fig. 4 illustrates the reproducibility of the state topographies over repeated data runs for one PD patient pre-music-cued exercise program. While the training converged to 9 states, different initializations resulted in two state types: persistent states that occurred in all runs, and transient, and less frequent states that varied between runs. Persistent states accounted for 70% of the FO. Consistency in converging to 9 states does not guarantee the states are identical due to the short data sequence and noise. However, there are states that persist, are reproducible, and stable between runs, while others seem to be run-specific. When more data were included by concatenating the participants of both groups, the resulting states were close to identical across repeated runs (Supp. Fig. 2).

**Fig. 4.**
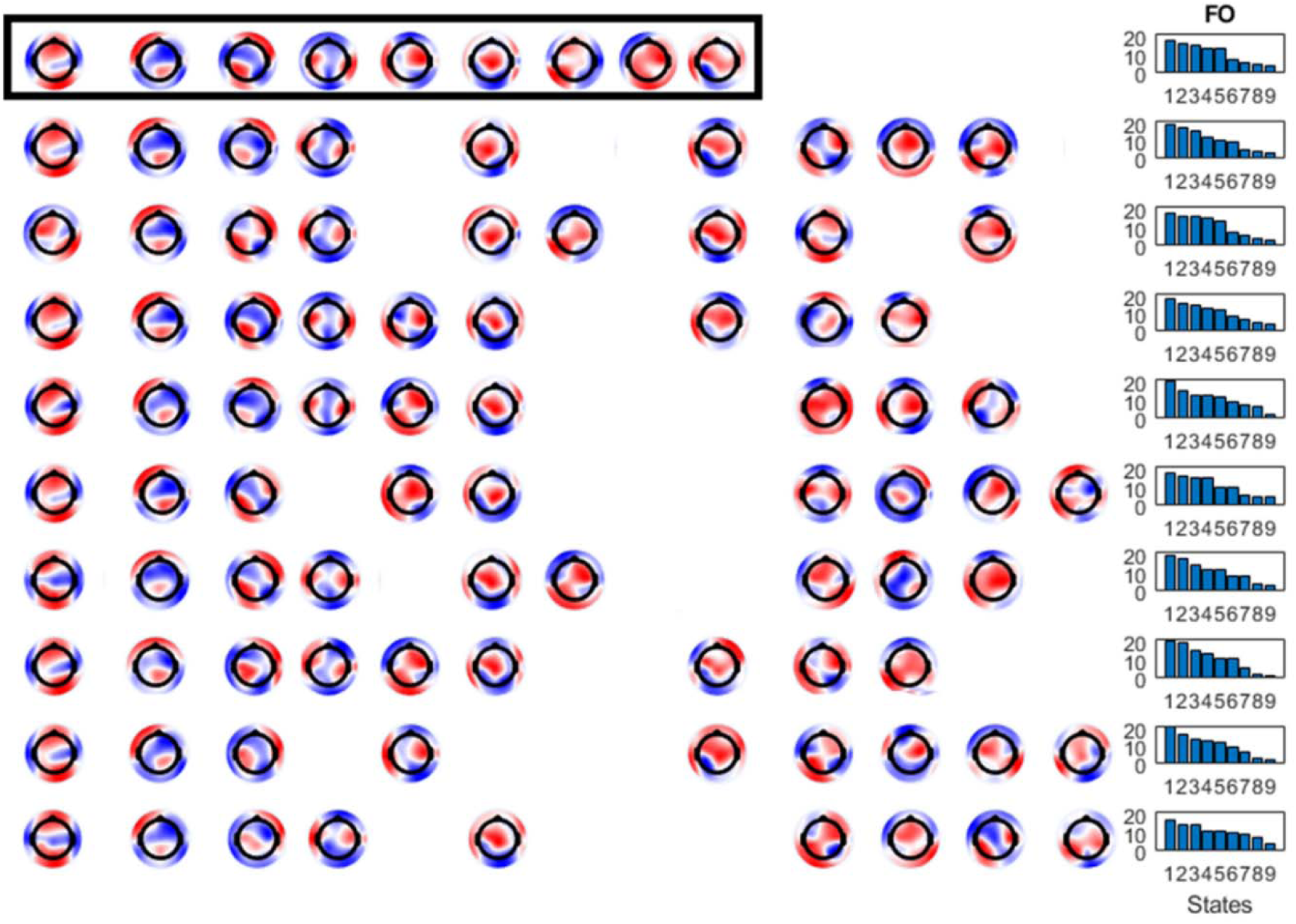
Reproducibility of Brain-State allocation in one individual. State topographies from ten repeated runs modeling 2 min of RS-EEG. The order of the state maps is arbitrary. Here we ordered the state maps using clustering and distance measures. Consistency in converging to 9 states, does not guarantee the states are identical, due to the short data sequence and noise. However, it is clear that there are states that persist, are reproducible, and stable between runs, while others seem to be run specific. Repeated runs of the same data help converge onto a set of consistent states, which also have the largest overlap in the fractional occupancy (FO) as shown on the **right column**, particularly when the sample size is small. The 4 most frequent states account for 70% of the observed data. Including more data, by concatenating the participants, results in convergence to consistent and reproducible states as shown in **Supp. Fig. 2**.

Second, the reproducibility of state maps was assessed for the group level data. After different independent runs for all data (2 groups), the best models were chosen according to the FE value. The models were trained on the concatenated data of 14 healthy control participants and 14 PD patients pre-music cued exercise program (total 28 data blocks). We used a fixed value of ***K*** = 9 states for data modeling. For all runs, the optimal model resulted in very similar state map patterns at the sensor level (see Supp. Fig. 2). Subsequently state sequences were used to extract individual metrics, revealing an individual’s fluctuations (and overlap) in resting state dynamics in the two groups. The FO range was computed for all participants for all the optimal models. Fig. 5 shows the resulting FO values and maps for each state and participant. Each point is the mean FO of one participant over 20 repeated runs, flanked by the standard deviation. An example of map sets for 10 different independent training model runs is shown in Supp. Fig. 2. Differences between maps were calculated. Maps belonging to different states showed statistical differences compared to maps belonging to the same state. The state maps for one model were used to analyze them for potential networks in the source space. The source analysis of the state maps is shown in Supp. Fig. 6.

**Fig. 5.**
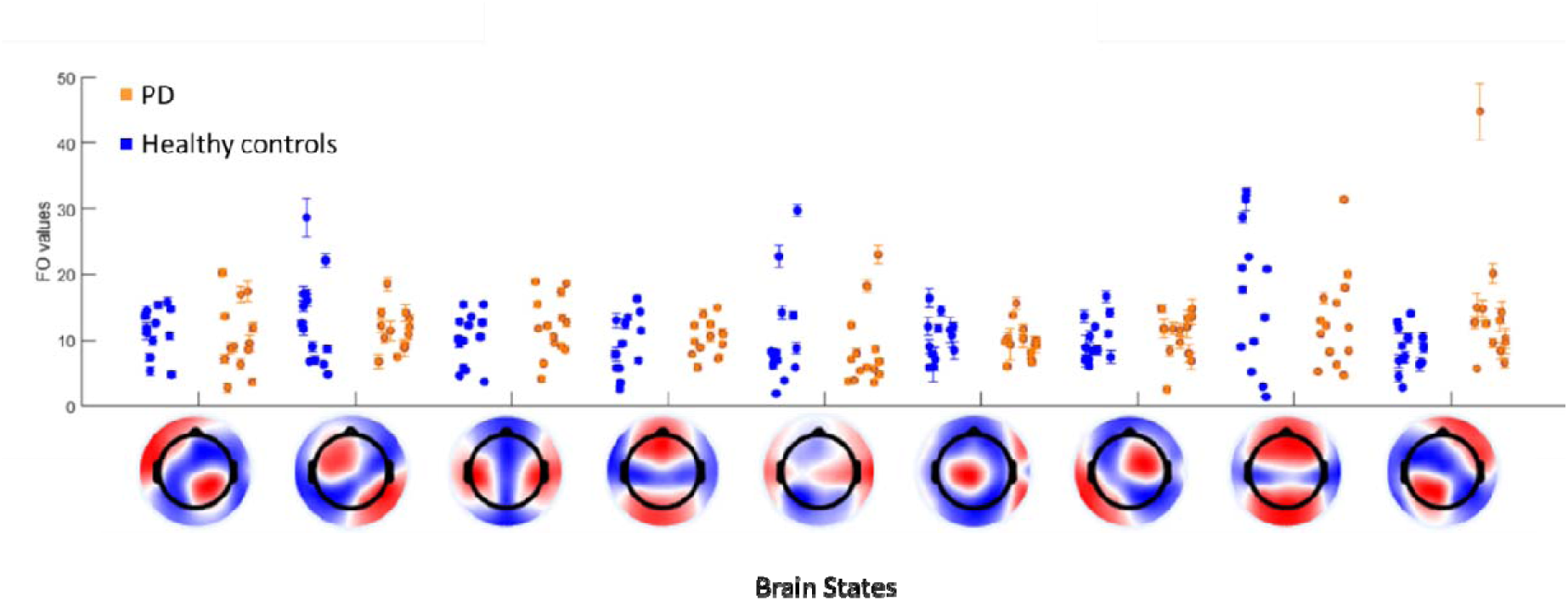
Individual differences and variation in fractional occupancy (FO) of the HsMM brain states, and their corresponding topographies, extracted from a group model of 2 min RS-EEG from 14 healthy control participants and 14 PD patients pre-exercise program. Data from both groups were concatenated to train 20 independent models. Subsequently state sequences were used to extract individual metrics, revealing the individual fluctuations (and overlap) in resting-state dynamics in the two groups. Each point is the mean FO of one participant over 20 repeated runs, flanked by the standard deviation. Consistency of FO over repeated modeling runs, averaged over participants within groups (healthy control participants versus PD patients pre-exercise program) is shown in **Supp. Fig. 3**. The equivalent figure showing Individual differences in dwell times (DT) is shown in **Supp. Fig. 5**.

Consistency of FO over repeated runs, averaged across participants within groups (healthy controls versus PD patients pre-exercise program) is shown in Supp. Fig. 3. The group differences for one of the model instantiations is shown in Supp. Fig. 4. The equivalent figure showing individual differences in DT is shown in Supp. Fig. 5.

### Exp. 3. Classification of individuals based on the dynamical features

The variability of classification across runs using the states’ FO showed the consistency of classifications of healthy participants and PD patients pre-exercise program for independent runs (Fig. 6 **top**). Secondly, we trained another classifier using the DTs as classification features (Fig. 6 **bottom**) with similar results to the classification of FO values. Classification scores did not show significant variance. While the figure highlights the large inter-subject variability, it also demonstrates the consistency of the dynamical features as representing individual characteristics. Indexes of participants and PD patients on the top graph are the same as in the bottom one. The order of the participants on the x-axis is arbitrary for ease of visualization. The classification of healthy control participants and PD patients pre-exercise program for each different set of features is shown in Supp. Fig. 8. Additionally, these classifications were compared to the classification using alpha frequency band values as features. The global alpha features for participants and PD patients are shown in Supp. Fig. 7. ROC curves and AUC comparing this classification to a classifier using as features the standard spectral analysis in the alpha band is shown in Supp. Fig. 9.

**Fig. 6.**
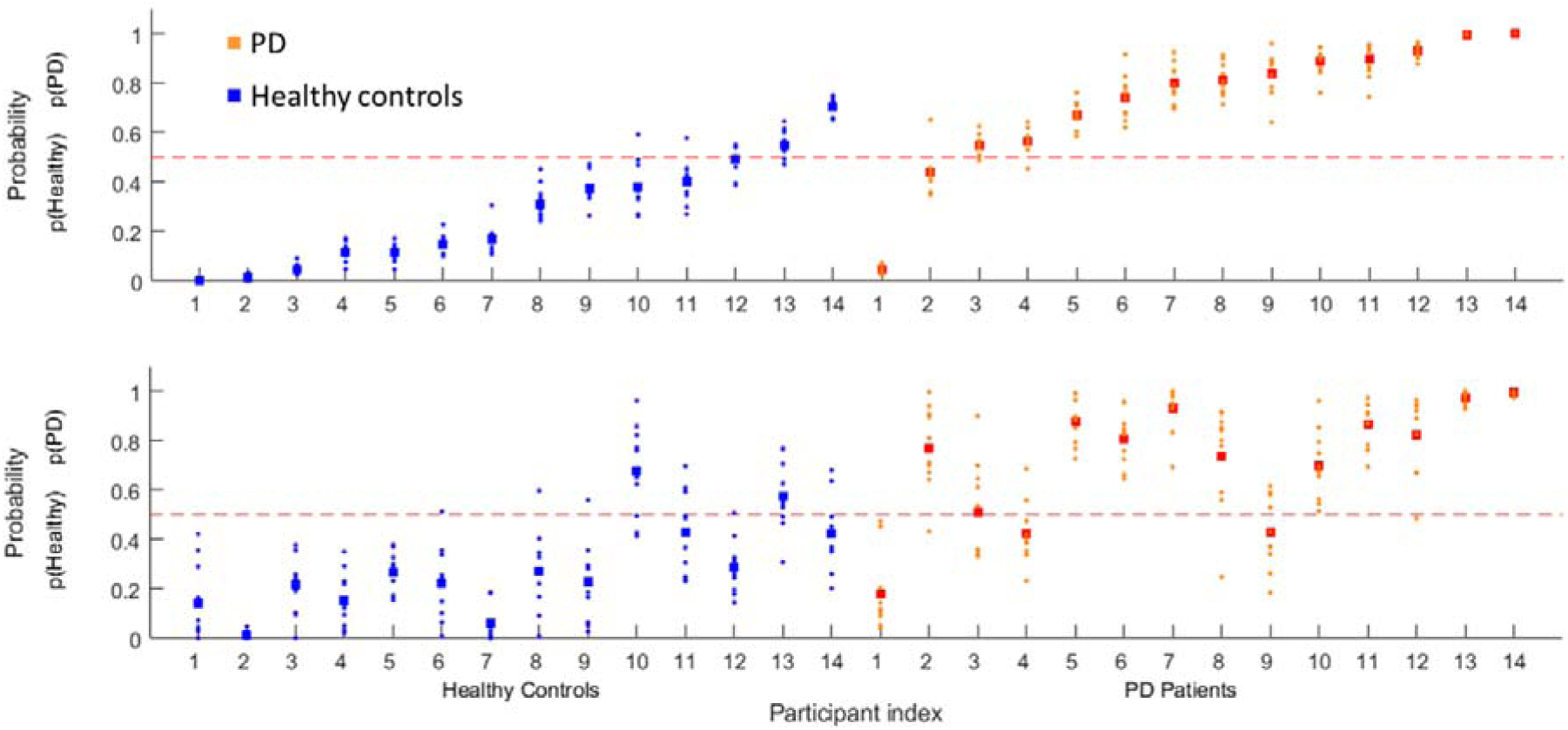
Classification of healthy control participants versus PD patients pre-exercise program based on the HsMM state metrics as dynamical features. Fractional occupancy (FO) (**top**), and dwell times (DT) (**bottom**) were extracted from the model in Fig. 5. The classifier was based on simple a Log-Regression which uses a General Linear Model (GLM) approach. Classifiers were constructed from **20** repeated HsMM modeling instantiations. Small circular points indicate the probability of each participant for each independent run. Square points indicate the average probability of each participant over the 20 runs. The dashed horizonal read line marks the 50% probability of being classified healthy versus PD. While the figure highlights the large intersubjective variability, it also demonstrates the consistency of the dynamical features as representing individual characteristics. Indexes of participants on the top graph are the same than in the bottom one. The order of the participants on the x-axis is arbitrary for ease of visualization. The classification of healthy control participants and PD patients pre-exercise program for each different set of features is shown in **Supp. Fig. 8.** ROC curves and AUC comparing this classification to a classifier using the standard spectral analysis in the alpha band as features is shown in **Supp. Fig. 9.**

### Exp. 4. Individual difference in response to music-cued exercise program

We used the trained classifiers and obtained HsMM features from the PD data pre- and post- music-cued movement exercise program. This is an indirect way to compare dynamic brain activity pattern changes after the exercise program. The classification of PD patients pre- and post-exercise program showed that some PD patients changed their HsMM patterns for FO and DTs. Fig. 7 shows the results of the classification of PD pre-exercise and post-exercise using FO (top) or DT (bottom) features. The figure shows the sensitivity of the dynamical features in response to the exercise program: in nearly 60% of the PD patients, the FO changed toward the characteristics of the healthy controls (blue dashed lines) after the exercise program, while 15% continued to progress in PD (blue dashed lines), and 25% remained unchanged (black dashed lines). The classification variations did not confirm significant differences for PD patients in each group. The classification and the alpha-band values as features are presented in the *Supplementary Material*.

**Fig. 7.**
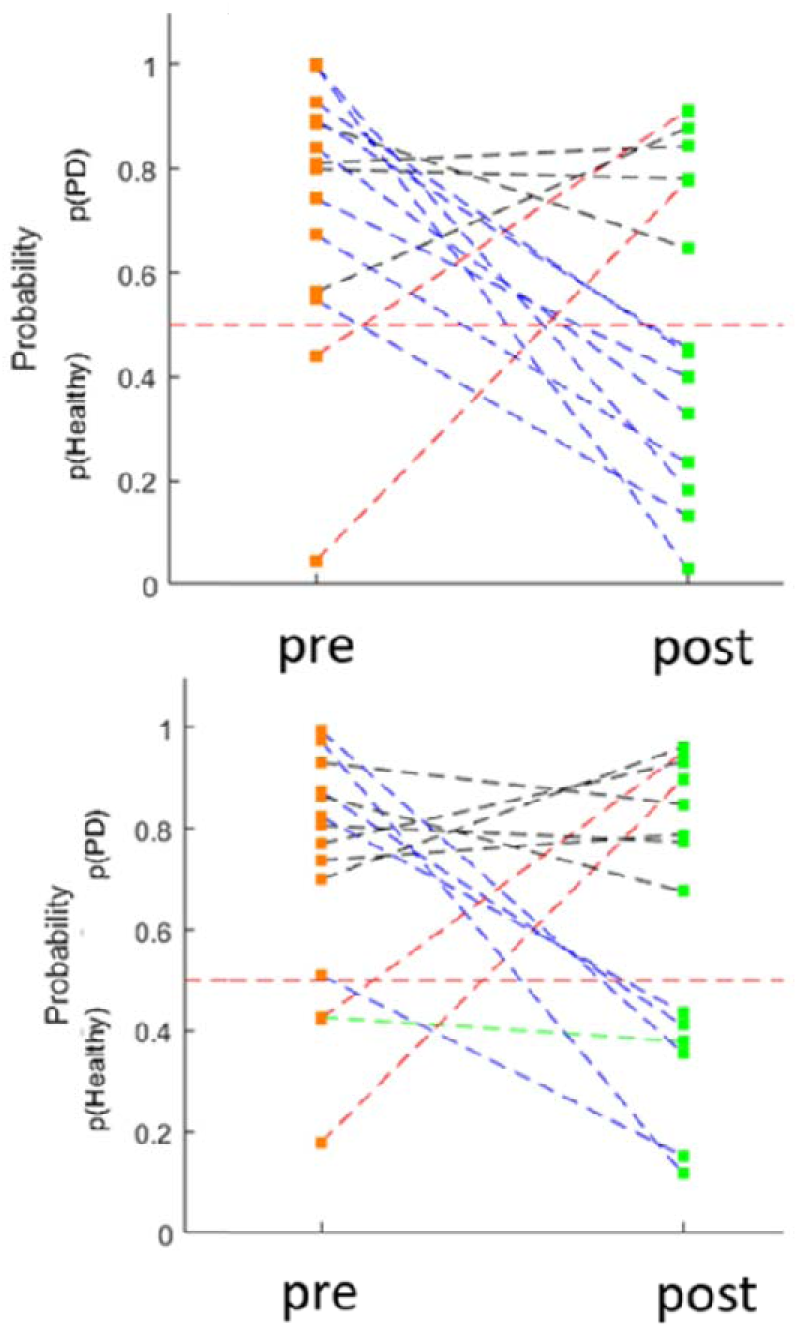
Change in classification score based on dynamical features of the PD patients post-exercise program. PD patients post-exercise program data were not part of the training of the HsMM model or the classifier. Instead, the state sequence was inferred from the model trained on the healthy control participants and PD patients pre-exercise program. After, the PD patients post-exercise program state sequences were used to obtain fractional occupancies (FO) and dwell times (DT). Classification was performed by using same classifier than in Fig. 6. using FO **Top**, or DT **Bottom.** The figure demonstrates the sensitivity of the dynamical features to the exercise program: nearly 60% of the PD patients had their FO moving towards the characteristics of the healthy controls (blue dashed lines) after the exercise program, while 15% become more like PD (blue dashed lines), and 25% remained unchanged (black dashed lines). FO was more sensitive to the exercise program than DT, suggesting that dynamical features could be used to reveal the specific effects interventions have on brain dynamics, and thus understand their mechanisms.

## 4. Discussion

The current study was motivated by modeling observable, spontaneous resting state (RS) changes in a clinical (PD) and healthy control group, considering that early RS dynamics changes can be useful markers of brain disease, ahead of measurable structural and functional change (Babiloni et al., 2013; Hatz et al., 2015). Specifically, cognitive changes might be related to such altered brain dynamics as seen in neurodegenerative diseases and correlated with early neurological and/or neuropsychiatric symptoms (Aarsland et al., 2009). Here we provide a proof of concept for the application of a class of dynamical brain-state allocation methods, namely Hidden semi-Markov Models (HsMM), that allow modeling clinical and non-clinical RS-EEG data. We modeled 2 min of spontaneous, eyes closed RS-EEG data from persons with PD and their matched healthy controls. We verified that the dynamical features, estimated by the model, are consistent. Moreover, these dynamical features allowed distinguishing persons with and without PD as well as changes (pre-/post-exercise program) induced by a music-cued exercise program that persons with PD underwent to improve their gait kinematics. These distinctions were not achieved by a standard power spectral alpha band data analysis (see Supp. Fig. 9). We report results indicative of the modeling robustness and reproducibility, particularly in relation to large intra- and inter-individual variability in RS-EEG and the stochasticity of the model estimation algorithms.

### 4.1 Generative models

Purely data-driven analysis methods such as classical classifiers, clustering, or principal components assume no knowledge about how measured data are generated, and thus cannot predict new data. Generative models, such as HMM or Kalman filters, describe a process that explains how observable data is created (Murphy, 2002). The model parameters are estimated from the data based on how well these parameters can recreate the data (Hsu et al., 2012). HMM/HsMM models assume a structure whereby a discrete hidden brain-state emits, for example, continuous observable EEG signals. The assumption is that the dynamics of the brain states, e.g., how long the brain takes in transitioning between states and resides in each state, conveys information about the brain’s functional state at a given time point. The physiological interpretation of the BSs is an ongoing research question. BSs seem to associate with a mixture of RSNs. Here, we used ELORETA to associate the EEG topographies from the BSs with the resting-state networks (RSNs). While the BSs to RSNs mapping is not one to one - rather, BSs seem to associate with a mixture of RSNs (see Supp. Fig. 6) - it supports the position that BS are operation modes in the brain that drive information flow and cognitive processes (Chen et al., 2019). Direct mapping between BS and RSNs has also been suggested, though it requires reliable estimates of the sources beforehand (Hernandez et al., 2022; Fukushima et al, 2018).

The estimation of the model parameters requires the optimization of a function, starting from a random initiation point. Given the complexity of the parameter space, there is no guarantee that every time the model is estimated, the optimization algorithm will converge to the same values. This raises practical questions about how reliable such models are and how meaningfully they can inform about dynamic changes in clinical profiles.

On the other hand, recent studies have used whole brain models to model brain activity and fit parameters to represent properties in empirical data, obtaining emergent properties. For example, large scale biophysical network models show metastability (Deco et al., 2017; Roberts et al., 2019). These types of models are structured, and their parameters are chosen to fit global features seen in biology, for instance, the global coupling (Castro et al., 2020). A generative model, like HsMM used here, is an approach in the middle of both mathematical models and empirical data, because it is a data driven approach that could offer a better explanation for some characteristics and properties in the data.

In the case of empirical data, clinical studies have shown variation in the characteristics of the brain’s resting state i.e., how they appear or transit (see for a comprehensive review Honcamp et al., 2022). An open question remains as to how this metastability or their features in the data correlate with symptoms or clinical severity or treatment response. Here we showed a feature that codifies a change or none after a non-pharmacological music-cued movement exercise program. This approach could be extended to other neurodegenerative conditions or to scenarios where dynamical features change in relation to experimental conditions or even to cognitive functions.

### 4.2 Considerations and recommendations for dynamical brain-state modeling

Several aspects must be considered for a useful HsMM implementation in clinical data. First, the data processing steps, and subsequent feature selection are critical factors for modeling. Data cleaning and transformation procedures depend on which feature will be modeled. For example, remaining data artifacts or noise could be modeled as an independent state by mistake. The selection of features, in this case the alpha band amplitude, determines what will be modeled and what conclusions can be drawn regarding the brain’s network dynamics. Second, due to the random naturalness of the model initialization, several runs/instantiations must be implemented to achieve reproducible and consistent results. This goes hand in hand with the amount of available data. Data sets with enough data points should be used to ensure consistent reproducibility of a model, which suggests that larger data sets offer better consistency and reproducibility. Third, regarding the number of brain states, there are many modeling approaches that use data driven criteria or rely on previously reported data. Here we used a data driven approach, taking advantage of a Bayesian implementation of the HsMM. We tested how the FE behaves across several data chunks to select the optimal number of states value. The navigation and search for the optimal number of states can be done for all data. However, this is extremely inefficient and computationally expensive. Therefore, we suggest using a random sample approach for choosing data chunks and searching the optimal number of states.

Here we also showed that we could model multigroup data without losing intra- and inter-individual differences. This is done by assuming a generative process common to all individuals, i.e., the same states and same state transitions, where each individual has a specific trajectory within this state space. This is a reasonable assumption, given the evidence that resting, task free, data, are manifestations of the underlying RSN dynamics, and that individual differences concern the variation on the utilization of those networks (Ceccheti et al., 2021).

The generative model approach goes beyond the phenomenological description, e.g., as in Microstates analysis (Michel and Koenig, 2018), to characterize the probabilistic process. This should be done understanding what aspects of the dynamics would need to be altered by interventions, such as brain stimulation techniques.

### 4.3 Dynamical brain-states in Parkinson’s Disease

We demonstrated the utility of the generative model approach using RS-EEG data from PD patients before and after a music-cued movement exercise program. Several studies have used Quantitative EEG (QEEG) analysis of the ongoing EEG spectrum to reveal alterations in multiple frequencies in PD (Babiloni et al., 2011; Cozac et al., 2016; Fonseca et al., 2015; Soikkeli et al., 1991; Stoffers et al., 2008; Mei et al., 2021); for example, the EEG signals of PD patients with dementia are distinctly slower than those of patients without dementia. Alpha power has been a powerful measure sensitive to cognitive impairment changes within PD (Babiloni et al., 2011). More recent efforts have focused on the classification of PD based on spectral features (Chu et al., 2021; Mei et al., 2021; Yang and Huang, 2022) correlated with neuropsychological tests (Jaramillo-Jimenez et al., 2021).

Here we showed that a simple classifier based on the dynamical features extracted from the HsMM outperforms classification based on the static spectral features. While the clinical significance of these results is outside the scope of the current work, it illustrates the power and utility of this modeling approach to sensitively monitor intra-and interindividual differences in clinical populations and beyond. This improved sensitivity should also reflect better correlations with neuropsychological tests and or clinical symptoms as suggested by Honcamp et al., (2022).

### 4.6 Limitations and future directions

We demonstrated a clear utility of the generative model approach to characterizing individual brain state variations. We motivated the analysis of the alpha band based on prior literature and characterized the dynamics at the level of the EEG electrodes. Future studies should generalize the current results by means of quantitative studies with larger participant samples and consider other frequency bands. Operating at the level of the EEG electrodes makes the task of interpreting the BSs physiologically harder. We linked the states to the estimation of the sources and showed that they map to a mixture of RSNs. A direct estimate of the RSNs at the source level would be more powerful, as proposed by Hernandez et al., (2022).

The current results show consistency and reliability with certain parameters and fixed values, e.g., the fixed number of states for group data modeling. This could be different in different settings and studies. We recommend using a similar approach to explore settings and parameters that better suit the data in a hypothesis-driven manner. We further suggest that the implementation used here offers a data-driven way for searching an optimal model using the Variational Bayes approach based on the FE for model comparisons. This has the advantage of resulting in distributions for the parameters instead of values. Future work could use this distribution characterization and relate it to behavioral data or task-dependent cognitive states in diverse populations.

### 4.7 Conclusion

The HsMM based characterization tool allowed describing and exploiting the information of brain dynamics obtained in a clinical group and healthy controls in a consistent way. PD patients’ brain dynamics showed differences for specific BS’s as compared to healthy participants brain dynamics. This was reflected in the HsMM metrics and in the classification of these metrics as features. Moreover, we showed that HsMM outcomes are consistent and replicable by assessing the variability of the values across different model instantiations.

Obtained BSs patterns showed differences between PD patients and healthy participants. Additionally, the clinical response to the music cued exercise program in PD patients was characterized by changes in brain dynamics profiles reflected by the HsMM models outputs and metrics.

Overall, HsMM is a powerful tool for analyzing brain activity dynamics and identifying hidden states that correspond to different BSs. HsMM can be used to analyze data from a variety of brain imaging techniques, and they have the potential to provide insights into the underlying mechanisms of brain function and the relationships between different brain states on a sub-second time scale.

### Significance

This work demonstrates that HsMM is a well-suited tool to describe brain dynamics obtained via RS-EEG. Moreover, this tool highlighted RS-EEG markers for persons with or without PD and changes in brain activity for some PD patients after an exercise program. In the future, the characterization of a disease and its response to any form of intervention might inform early diagnosis before symptoms occur.

## Data and code availability

Data will be made available via a request to the authors and pending review. A data sharing agreement may be required before access is granted. The software used in this work is available on https://github.com/daraya78/BSD.

## Credit authorship contribution statement

Sonja A. Kotz: Conceptualization, recruitment and data collection, and manuscript writing and editing.

Aland Astudillo: data analysis, modeling, and manuscript writing and editing.

David Araya: Software implementation.

Simon Dalla Bella: Recruitment and data collection.

Nelson Trujillo-Barreto: Software implementation, review & editing.

Wael El-Deredy: Conceptualization and manuscript writing and editing.

## Acknowledgments

We are grateful to advice from Patricio Orio, Lucía Zepeda, Julio Rodiño, Grace Whitaker and, Hanna Honcamp for discussions and assistance.

