## Supplemental Material for "Modeling EEG Resting-State Brain Dynamics: A proof of concept for clinical studies"

**Title**

### **1. Choosing the optimal number of states**

One important aspect of the Brain-States (BSs) allocation is the selection of the number of states. In order to select an optimal number of states, we used a data-driven approach. The implementation used here utilizes a variational approach which uses a Free-Energy (FE) criteria for finding the optimal model (Trujillo-Barreto et al., 2019). We tested how the FE behaves for different training runs and potentially changing the number of states. This is not practical for the entire data set (i.e., modeling all healthy control participants and PD patients). Instead, we tested the convergence of the model for different independent blocks of transformed RS-EEG data.

We tested the model convergence to a consistent number of states over repeated runs. Data from a single participant was used for training 50 models, while changing the number of states from 4 to 10. The FE was used as a model comparison/model selection criterion. The best model, with the largest FE, corresponds to the optimal number of states (see Fig. 3). Data for each participant was used for training models and to verify the convergence for each one independently. We repeated this experiment over ten randomly chosen participants to estimate the optimal number of states.

Supp. Fig. 1 shows the reproducibility of BSs allocation in single participant level. The figure shows the convergence according to the FE to a number of states (***K*** = 9) over repeated runs for individuals only. Supp. Fig. 1 **top left** graph shows the different obtained FE values across repetitions of the model. Supp. Fig. 1 **top right** graph shows how the FE values change according to the number of states values explored. Supp. Fig. 1 **bottom left** graph shows how the FE values change through iterations for the optimal model. Supp. Fig. 1 **bottom right** graph shows how the FE values change through iterations for all the models including the non-optimal ones. The most common value was ***K*** = 9 states, which was then applied to subsequent group experiments. This approach serves as an example to further studies in similar or different settings.

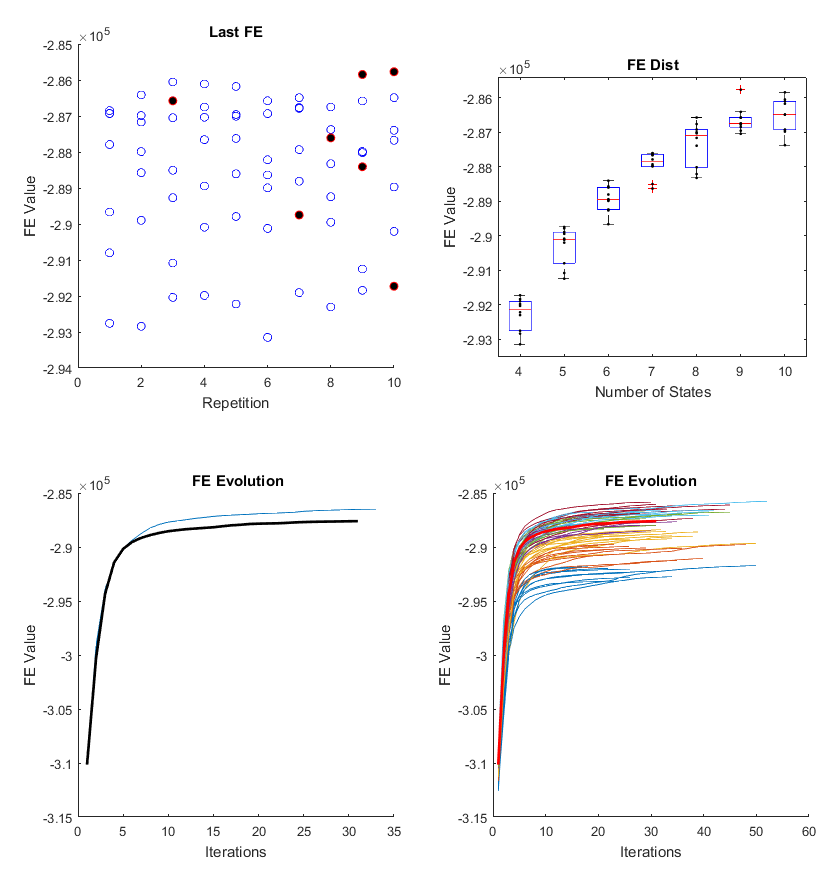

**Supp. Fig. 1.** Reproducibility of Brain-States (BSs) allocation in single participant level. The figure shows the convergence according to the Free-Energy (FE) to a number of states (***K*** = 9) over repeated runs for individuals only. Data for each participant was used for training models and to verify the convergence for each one independently. The training navigates the number of states from 4 to 10 and it determines the most probable number of states. We repeated this experiment over ten randomly chosen participants to estimate the number of states, which was then applied to subsequent group experiments. The most common value was ***K*** = 9 States. **Top left** graph shows the different obtained FE values across repetitions of the model. **Top right** graph shows how the FE values change according to the number of states values explored. **Bottom left graph** shows how the FE values change through iterations for the optimal model. **Bottom right graph** shows how the FE values change through iterations for all the models including the non-optimal ones.

### **2. Variability of estimated dynamical features over repeated runs**

In order to test the consistency and reproducibility of the model outcomes, we performed both modeling at the level of single participant and at the group level. Here we present the additional results at the group level.

*Variability across repeated runs at state maps*

The variability across repeated independent runs was tested. We trained 20 independent models using the concatenated alpha-band amplitude envelope data from 14 healthy participants and 14 PD patients pre-exercise program (total 28 data blocks), and then we obtained state maps, state sequences, and metrics for each participant/patient. Because the order of the state is arbitrary each time that the model finishes the training, we identified the similarity between the state maps using clustering and distance measures. We use a combination of Euclidian distance and clustering in a Principal Component Analysis (PCA) subspace of the spatial information of maps to match maps from different model instantiations to resort them. Information of this decomposition and resorting is not shown here.

*Variability across repeated runs at fractional occupancy (FO)*

The variability of individual mean fractional occupancies (FO) for 20 models was quantified. We trained 20 independent models using the data from 14 healthy control participants and 14 PD patients pre-exercise program. For each model we obtained FO metrics for each participant. The values across participants (from Fig. 5) were averaged for each state/group. States were sorted according to the mean FO in descending order (for healthy participants).

*Variability across individuals for one model instantiation at FO*

We assessed the variability of HsMM metrics FO values for one model. One of the 20 model instantiations was assessed (group model). Subsequently state sequences were used to extract individual metrics, revealing the individual fluctuations (and overlap) in resting-state dynamics in the two groups. Here we show the individual differences in FO per state.

*Variability across repeated runs at dwell times (DT)*

Dwell times were fitted using a log-normal obtaining two parameters (mu and sigma) for each participant in each state and model instantiation. We computed individual differences in DT for mu and sigma per state, extracted from a group model of 2 min RS-EEG from 14 healthy control participants and 14 PD patients pre-exercise program concatenated data (total 28 data blocks). Subsequently state sequences were used to extract individual metrics, revealing the individual fluctuations (and overlap) in resting-state dynamics in the two groups. Fig. 5 shows individual differences in FO**.**

Supp. Fig. 2 shows the reproducibility of brain state allocation in group modeling and the variability of states maps. Ten repetitions are shown modeling group data. The repeated runs converged to 9 states that are very similar. Distances between the maps show a small variance (not shown here).

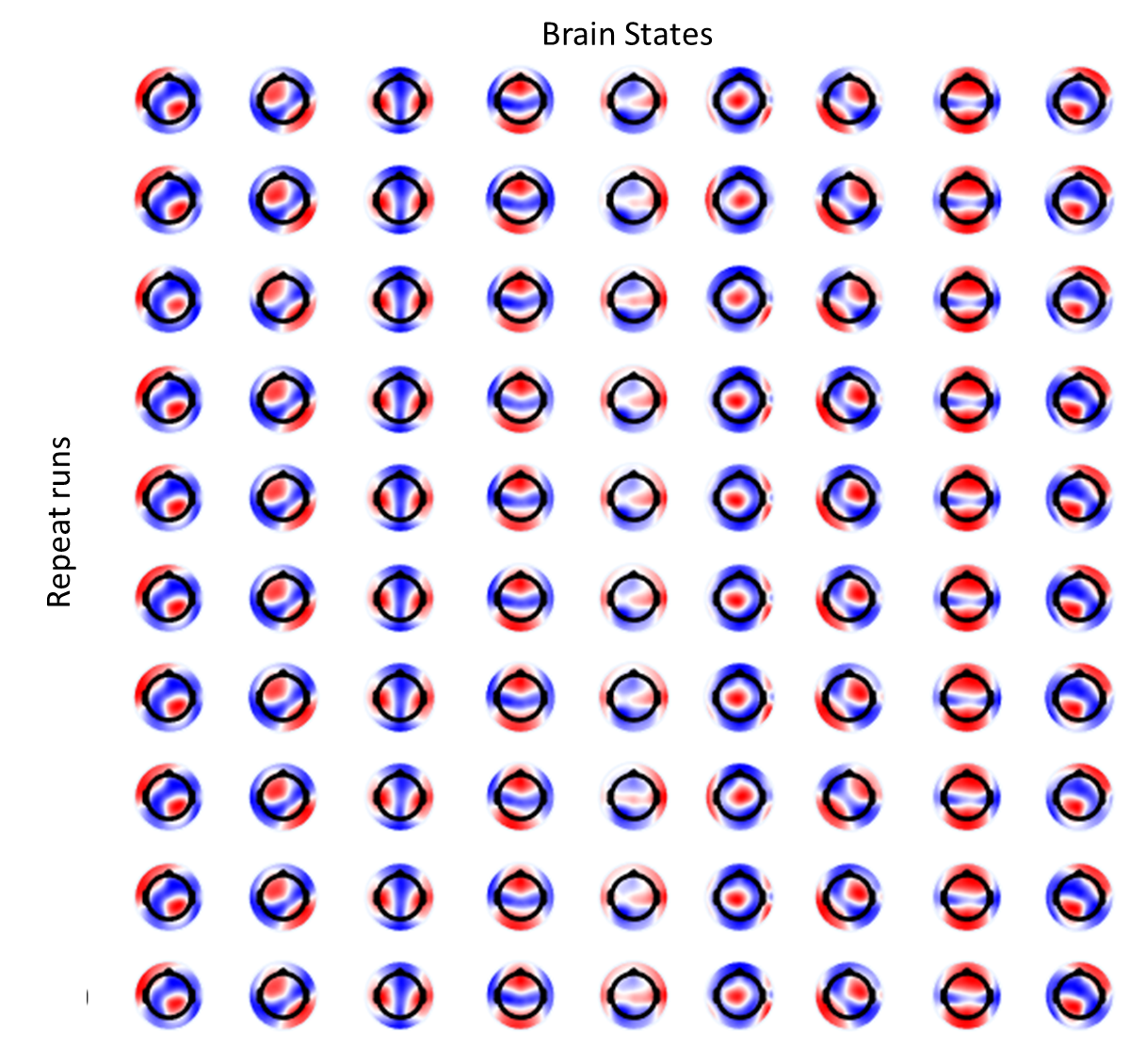

**Supp. Fig. 2**. Reproducibility of brain state allocation in group modeling. Ten repetitions are shown modeling data from 14 healthy control participants and 14 PD patients pre-exercise program concatenated with 2 min of RS-EEG each (total 28 data blocks). Because the order of the state maps is arbitrary each time that the model finish the training, we identified the similarity between the state maps using clustering and distance measures. The repeated runs converged to 9 states that are very similar.

Supp. Fig. 3 shows the consistency of FO over repeat modeling runs, averaged over participants for each group. Each point represents a group of 20 agglutinated points showing how low is the variance across models.

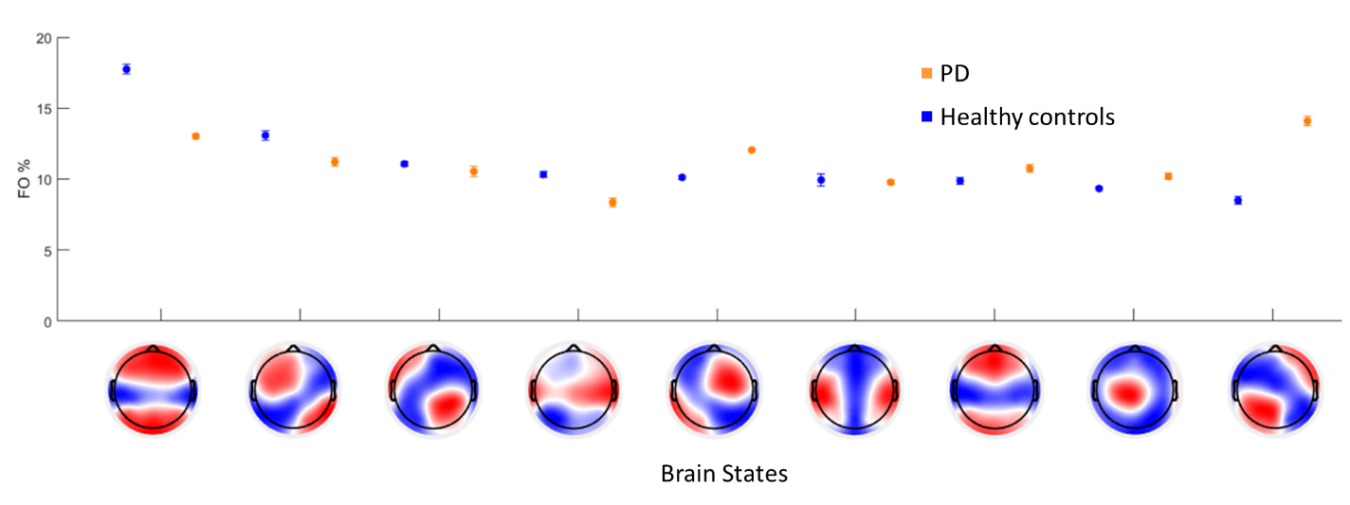

**Supp. Fig. 3**. Consistency of fractional occupancy (FO) over repeated modeling runs, averaged over participants for each group. Variability of individual mean FO for 20 models was quantified. We trained 20 independent models using the data from 14 healthy control participants and 14 PD patients pre-exercise program. For each model we obtained FO metrics for each participant. The values across participants (from **Fig. 5**) were averaged for each state/group. States were sorted according to the mean FO in descending order (for healthy participants). Thus, each point here is group of 20 agglutinated points showing how low is the variance across models.

Supp. Fig. 4 shows the group variability of HsMM metrics FO values for 1 model instantiation. Each point corresponds to the FO value for each participant/patient in each state. The error bars show the variance across participants for each state. Solid circular markers indicate the group mean for each state. States were sorted according to the mean FO in descending order (for healthy participants).

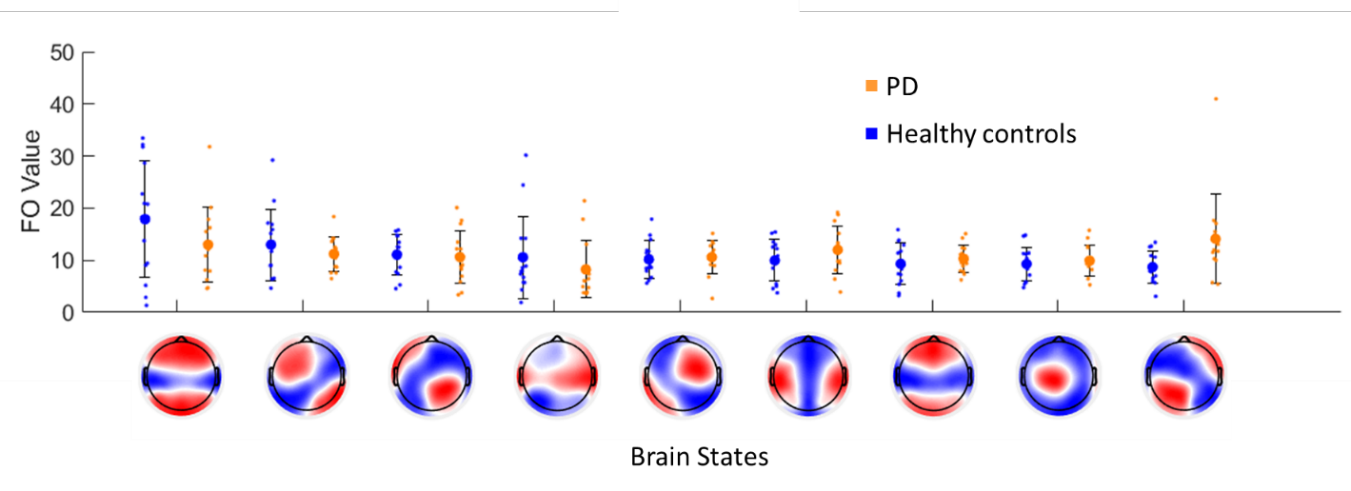

**Supp. Fig. 4.** Group variability of HsMM metrics values for one model. Here we show the individual differences in fractional occupancy (FO) per state, extracted from a group model of 2 min RS-EEG from 14 healthy control participants and 14 PD patients pre-exercise program (total 28 data blocks). Subsequently state sequences were used to extract individual metrics, revealing the individual fluctuations (and overlap) in resting-state dynamics in the two groups. Each point corresponds to the FO value for each participant/patient in each state. The error bars show the variance across participants for each state. Solid circular markers indicate the group mean for each state. States were sorted according to the mean FO in descending order (for healthy participants).

Supp. Fig. 5 shows the individual differences in DT for mu and sigma per state, obtained from a group model. While there is no significant difference between the groups, in general, DTs are longer and more variable in the healthy group. States were sorted according to the mean FO in descending order (for healthy participants). We could observe that the range of variability for HsMM metrics FO and DT are acceptable and that for different model instantiations which start from random initializations, the final results are suitable to be consistent and replicable.

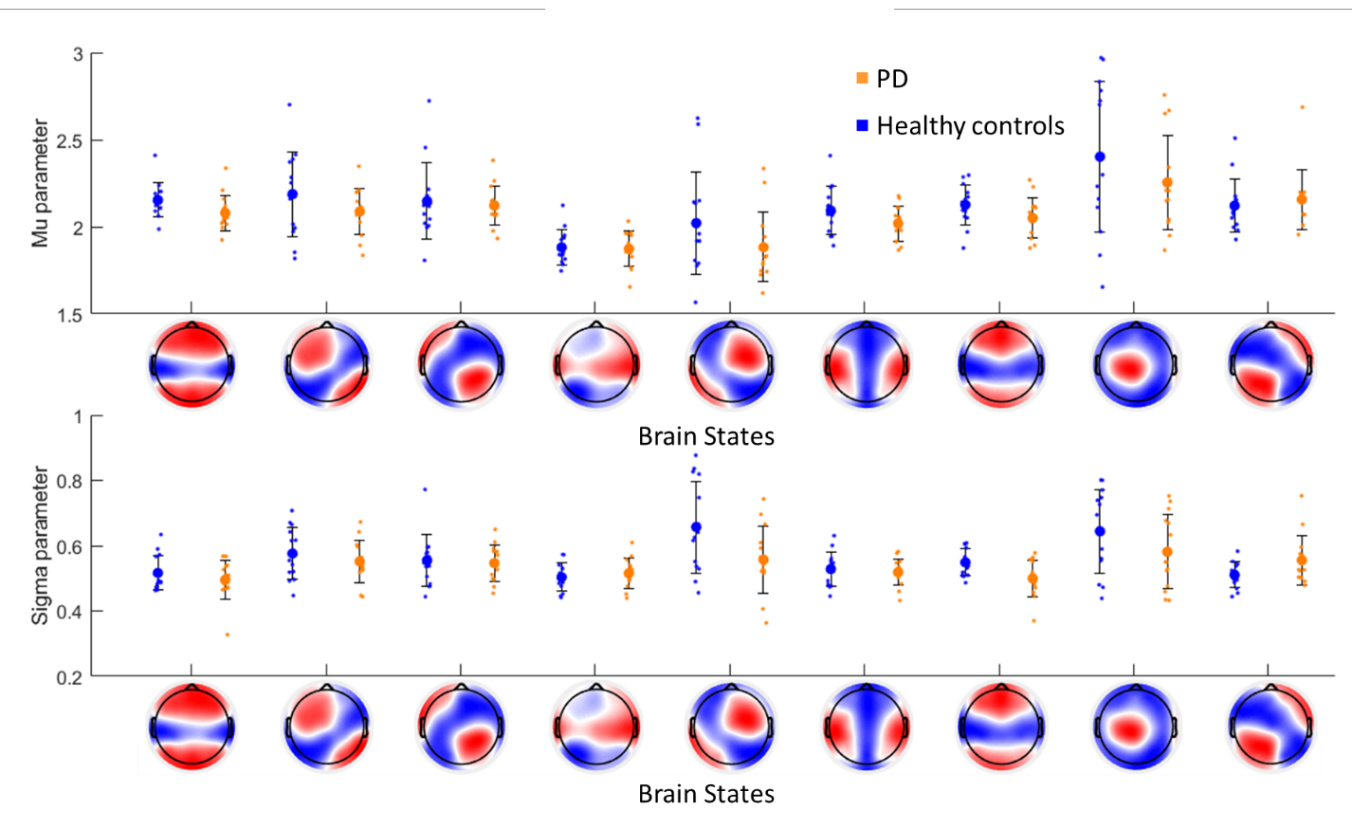

**Supp. Fig. 5.** Individual differences in dwell time (DT) mean and variance per state, extracted from a group model of 2 min RS-EEG from 14 healthy control participants and 14 PD patients pre-exercise program concatenated data (total 28 data blocks). Subsequently state sequences were used to extract individual metrics, revealing the individual fluctuations (and overlap) in resting-state dynamics in the two groups. The empirical state dwell times were fitted as a log-normal curve for each participant/state. Two parameters were obtained to characterize each dwell time distribution (mu and sigma). Equivalent figure showing Individual differences in fractional occupancy (FO) is shown in **Fig. 5**. While there is no significant difference between the groups, in general, DTs are longer and more variable in the healthy group. States were sorted according to the mean FO in descending order (for healthy participants).

### **3. Neural generators of Brain-State topographies**

We evaluated the potential neural generators of the brain state topographies obtained for the HsMM modeling. The goal is to verify if these HsMM states maps relate to known network configurations, in terms of cortical zones related to Resting State Networks (RSNs) to some extent.

State topographies were related to brain areas. We used the topographical information of each brain state to perform source reconstruction. The HsMM state maps were obtained for a model instantiation trained on concatenated alpha-band amplitude envelope data of 14 healthy control participants and 14 PD patients pre-exercise program (total 28 concatenated data blocks) (see Fig. 5). We used a source analysis method based on exact Low Resolution Electromagnetic Tomography Analysis (eLORETA) (Pascual-Marqui. 2007), by using LORETA-KEY v20170220 software (the Key Institute for Brain-Mind Research, Zurich, Switzerland). The analysis approach it was applied similarly to ICA based source analysis. In this approach, the ICA topography for each independent component serves as the input to the source reconstruction. Here, instead of using the ICA topography, we used the HsMM state topography as input to the source analysis for each state independently. After each source analysis, we obtained a set of possible sources for each HsMM state. We defined a threshold for each analysis to obtain the most probable active cortical structures related to each HsMM state activation.

Supp. Fig. 6 shows the estimated cortical generators of the HsMM brain states. Each state shows a slightly different pattern of activation at the level of the sources. The Supplementary Table 1 shows the cortical nodes for the source reconstruction of each map. The resulting cortical areas showed multiple overlapping structures when compared to classical resting state networks (RSNs). This may suggest that the relationship between HsMM states and RSNs are not one to one, which is in line with recent findings (Hunyadi et al., 2019).

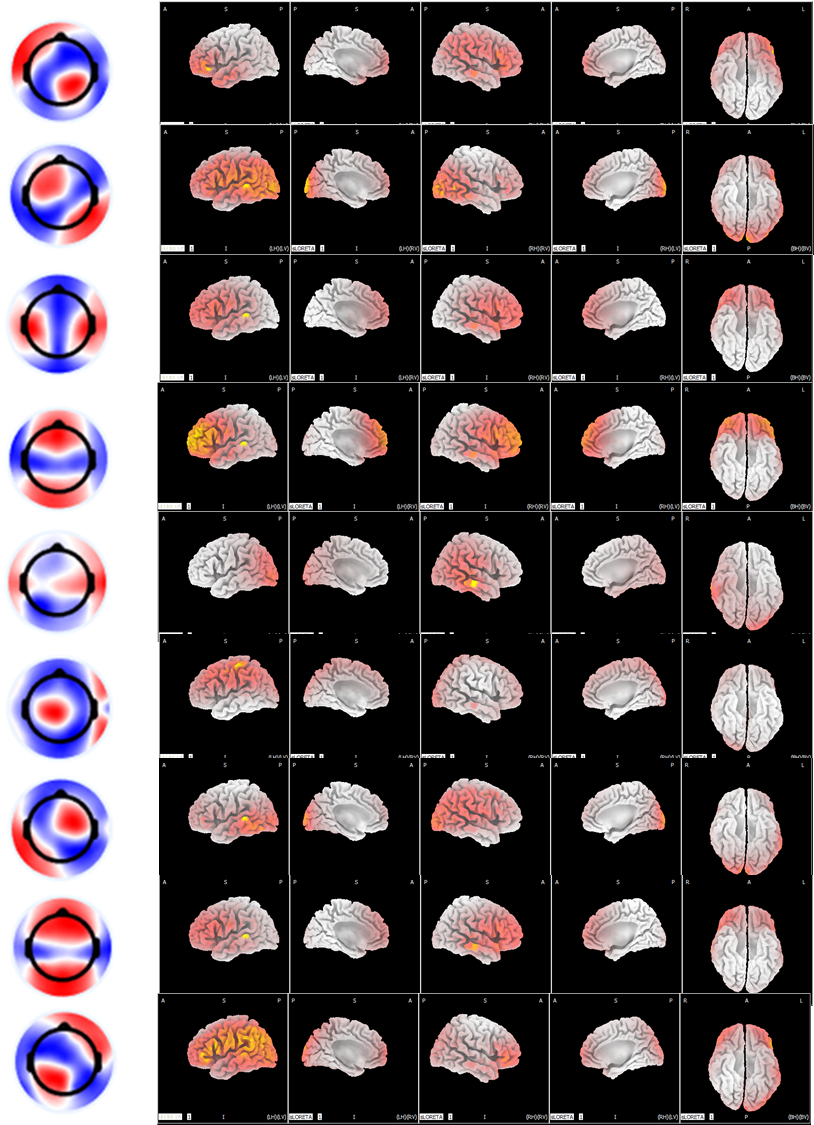

**Supp. Fig. 6.** Estimated cortical generators of the HsMM brain states. Brain states topographies were obtained for the multi group HsMM model of 14 healthy control participants and 14 PD patients pre-exercise program (**Fig. 5**). For each state map, the cortical sources were estimated using the exact low resolution electromagnetic tomography analysis (ELORETA) method (**Pascual-Marqui, 2007**). The resulting cortical areas showed multiple overlapping structures when comparing to classical resting state networks (RSNs), which may suggest that the relationship between HsMM states and RSNs are not one to one, which is line with recent findings (**Hunyadi et al., 2019**).

**Supplementary Table 1. Cortical nodes for source reconstruction of state maps**

| HsMM Brain State | N voxels | N structures | Structures |
| --- | --- | --- | --- |
| 1 | 431 | 19 | Frontal Lobe, Inferior Frontal Gyrus |
|  |  |  | Temporal Lobe, Middle Temporal Gyrus |
|  |  |  | Parietal Lobe, Supramarginal Gyrus |
|  |  |  | Frontal Lobe, Middle Frontal Gyrus |
|  |  |  | Temporal Lobe, Superior Temporal Gyrus |
|  |  |  | Temporal Lobe, Supramarginal Gyrus |
|  |  |  | Frontal Lobe, Superior Frontal Gyrus |
|  |  |  | Parietal Lobe, Inferior Parietal Lobule |
|  |  |  | Frontal Lobe, Precentral Gyrus |
|  |  |  | Parietal Lobe, Postcentral Gyrus |
|  |  |  | Temporal Lobe, Inferior Temporal Gyrus |
|  |  |  | Temporal Lobe, Angular Gyrus |
|  |  |  | Frontal Lobe, Sub-Gyral |
|  |  |  | Parietal Lobe, Superior Parietal Lobule |
|  |  |  | Frontal Lobe, Medial Frontal Gyrus |
|  |  |  | Frontal Lobe, * |
|  |  |  | Parietal Lobe, Angular Gyrus |
|  |  |  | Frontal Lobe, Orbital Gyrus |
|  |  |  | Frontal Lobe, Rectal Gyrus |
| 2 | 460 | 21 | Temporal Lobe, Middle Temporal Gyrus |
|  |  |  | Temporal Lobe, Superior Temporal Gyrus |
|  |  |  | Occipital Lobe, Middle Occipital Gyrus |
|  |  |  | Occipital Lobe, Cuneus |
|  |  |  | Temporal Lobe, Inferior Temporal Gyrus |
|  |  |  | Occipital Lobe, Inferior Occipital Gyrus |
|  |  |  | Occipital Lobe, Lingual Gyrus |
|  |  |  | Frontal Lobe, Precentral Gyrus |
|  |  |  | Occipital Lobe, Middle Temporal Gyrus |
|  |  |  | Frontal Lobe, Inferior Frontal Gyrus |
|  |  |  | Temporal Lobe, Transverse Temporal Gyrus |
|  |  |  | Parietal Lobe, Inferior Parietal Lobule |
|  |  |  | Parietal Lobe, Postcentral Gyrus |
|  |  |  | Parietal Lobe, Supramarginal Gyrus |
|  |  |  | Frontal Lobe, Middle Frontal Gyrus |
|  |  |  | Parietal Lobe, Angular Gyrus |
|  |  |  | Temporal Lobe, Angular Gyrus |
|  |  |  | Temporal Lobe, Supramarginal Gyrus |
|  |  |  | Occipital Lobe, Superior Occipital Gyrus |
|  |  |  | Parietal Lobe, Precuneus |
|  |  |  | Frontal Lobe, Superior Frontal Gyrus |
| 3 | 2544 | 26 | Temporal Lobe, Superior Temporal Gyrus |
|  |  |  | Frontal Lobe, Precentral Gyrus |
|  |  |  | Frontal Lobe, Inferior Frontal Gyrus |
|  |  |  | Temporal Lobe, Middle Temporal Gyrus |
|  |  |  | Frontal Lobe, Middle Frontal Gyrus |
|  |  |  | Frontal Lobe, Superior Frontal Gyrus |
|  |  |  | Frontal Lobe, Medial Frontal Gyrus |
|  |  |  | Parietal Lobe, Postcentral Gyrus |
|  |  |  | Frontal Lobe, Orbital Gyrus |
|  |  |  | Temporal Lobe, Transverse Temporal Gyru |
|  |  |  | Frontal Lobe, Sub-Gyral |
|  |  |  | Frontal Lobe, * |
|  |  |  | Parietal Lobe, Inferior Parietal Lobule |
|  |  |  | Limbic Lobe, Anterior Cingulate |
|  |  |  | Parietal Lobe, Supramarginal Gyrus |
|  |  |  | Sub-lobar, Insula |
|  |  |  | Frontal Lobe, Rectal Gyrus |
|  |  |  | Temporal Lobe, Supramarginal Gyrus |
|  |  |  | Sub-lobar, Extra-Nuclear |
|  |  |  | Limbic Lobe, Cingulate Gyrus |
|  |  |  | Temporal Lobe, Inferior Temporal Gyrus |
|  |  |  | Frontal Lobe, Extra-Nuclear |
|  |  |  | Frontal Lobe, Postcentral Gyrus |
|  |  |  | Frontal Lobe, Cingulate Gyrus |
|  |  |  | Frontal Lobe, Subcallosal Gyrus |
|  |  |  | Temporal Lobe, Inferior Frontal Gyrus |
| 4 | 46 | 6 | Temporal Lobe, Superior Temporal Gyrus |
|  |  |  | Frontal Lobe, Middle Frontal Gyrus |
|  |  |  | Frontal Lobe, Superior Frontal Gyrus |
|  |  |  | Frontal Lobe, Precentral Gyrus |
|  |  |  | Frontal Lobe, Inferior Frontal Gyrus |
|  |  |  | Frontal Lobe, Medial Frontal Gyrus |
| 5 | 140 | 16 | Temporal Lobe, Middle Temporal Gyrus |
|  |  |  | Occipital Lobe, Cuneus |
|  |  |  | Temporal Lobe, Superior Temporal Gyrus |
|  |  |  | Occipital Lobe, Middle Occipital Gyrus |
|  |  |  | Occipital Lobe, Inferior Occipital Gyrus |
|  |  |  | Occipital Lobe, Lingual Gyrus |
|  |  |  | Parietal Lobe, Supramarginal Gyrus |
|  |  |  | Temporal Lobe, Inferior Temporal Gyrus |
|  |  |  | Occipital Lobe, Fusiform Gyrus |
|  |  |  | Temporal Lobe, Angular Gyrus |
|  |  |  | Parietal Lobe, Postcentral Gyrus |
|  |  |  | Occipital Lobe, Middle Temporal Gyrus |
|  |  |  | Parietal Lobe, Inferior Parietal Lobule |
|  |  |  | Occipital Lobe, Superior Occipital Gyrus |
|  |  |  | Parietal Lobe, Precuneus |
|  |  |  | Temporal Lobe, Supramarginal Gyrus |
| 6 | 7 | 2 | Parietal Lobe, Postcentral Gyrus |
|  |  |  | Parietal Lobe, Inferior Parietal L |
| 7 | 439 | 21 | Temporal Lobe, Superior Temporal Gyrus |
|  |  |  | Temporal Lobe, Inferior Temporal Gyrus |
|  |  |  | Occipital Lobe, Middle Occipital Gyrus |
|  |  |  | Occipital Lobe, Cuneus |
|  |  |  | Occipital Lobe, Lingual Gyrus |
|  |  |  | Temporal Lobe, Middle Temporal Gyrus |
|  |  |  | Temporal Lobe, Angular Gyrus |
|  |  |  | Occipital Lobe, Inferior Occipital Gyrus |
|  |  |  | Occipital Lobe, Middle Temporal Gyrus |
|  |  |  | Temporal Lobe, Supramarginal Gyrus |
|  |  |  | Parietal Lobe, Supramarginal Gyrus |
|  |  |  | Frontal Lobe, Precentral Gyrus |
|  |  |  | Occipital Lobe, Superior Occipital Gyrus |
|  |  |  | Parietal Lobe, Angular Gyrus |
|  |  |  | Parietal Lobe, Inferior Parietal Lobule |
|  |  |  | Parietal Lobe, Precuneus |
|  |  |  | Frontal Lobe, Middle Frontal Gyrus |
|  |  |  | Parietal Lobe, Postcentral Gyrus |
|  |  |  | Frontal Lobe, Inferior Frontal Gyrus |
|  |  |  | Parietal Lobe, Superior Parietal Lobule |
|  |  |  | Temporal Lobe, Transverse Temporal Gyrus |
| 8 | 3 | 3 | Temporal Lobe, Superior Temporal Gyrus |
|  |  |  | Temporal Lobe, Middle Temporal Gyrus |
|  |  |  | Frontal Lobe, Inferior Frontal Gyrus |
| 9 | 659 | 22 | Frontal Lobe, Inferior Frontal Gyrus |
|  |  |  | Frontal Lobe, Precentral Gyrus |
|  |  |  | Parietal Lobe, Supramarginal Gyrus |
|  |  |  | Parietal Lobe, Inferior Parietal Lobule |
|  |  |  | Temporal Lobe, Middle Temporal Gyrus |
|  |  |  | Parietal Lobe, Angular Gyrus |
|  |  |  | Parietal Lobe, Postcentral Gyrus |
|  |  |  | Temporal Lobe, Supramarginal Gyrus |
|  |  |  | Temporal Lobe, Superior Temporal Gyrus |
|  |  |  | Temporal Lobe, Angular Gyrus |
|  |  |  | Occipital Lobe, Middle Occipital Gyrus |
|  |  |  | Occipital Lobe, Cuneus |
|  |  |  | Occipital Lobe, Superior Occipital Gyrus |
|  |  |  | Parietal Lobe, Precuneus |
|  |  |  | Frontal Lobe, Middle Frontal Gyrus |
|  |  |  | Parietal Lobe, Superior Parietal Lobule |
|  |  |  | Occipital Lobe, Middle Temporal Gyrus |
|  |  |  | Occipital Lobe, Inferior Occipital Gyrus |
|  |  |  | Temporal Lobe, Transverse Temporal Gyrus |
|  |  |  | Temporal Lobe, Inferior Temporal Gyrus |
|  |  |  | Sub-lobar, Insula |
|  |  |  | Frontal Lobe, Superior Frontal Gyrus |

### **4. Comparison between classifiers of dynamical features and spectral features**

In order to evaluate the potential of HsMM metrics as features to characterize brain activity dynamics, we compared them with commonly used alpha-band values for brain activity classification.

*Alpha power estimation*

We calculated the alpha band amplitude values (from now, alpha power) for each cleaned RS-EEG data set of healthy control participant and PD patient pre-exercise program. After data pre-processing, 2 min clean eyes closed RS-EEG data segments were extracted. We computed the power spectrum density (PSD) using the Welch method periodogram for each data set. PSD of clean segments were averaged across all channels and the peak of the alpha band (8-14 Hz) was used to get the amplitude of each data set for further analysis. The alpha power for 9 representative channels were used for each data set (F3, Fz, F4, C3, Cz, C4, P3, Pz, P4). These values were then used as features for classification.

*Classification of participants according to metrics*

We compared the classification performance between alpha power features (here the mean peak value of the alpha band for each data set) and the HsMM features (here the FO values, and states DT values). We used a simple log-regression classifier (LRC), which is based on the General Linear Model (GLM). We measured the general classification performance based on the Receiver Operator Characteristic (ROC) curve, and the ROC area under curve (AUC). For the HsMM features sets, we used 3 different metrics as inputs: FO and DT. In the case of DT, because the durations follow a log-normal distribution, we fitted a curve for each empirical duration value set obtaining two parameters: *mu* and *sigma*. DT *mu* and DT sigma were used as classification features. For spectral features, we used the alpha power for 9 representative channels for each participant/patient.

Supp. Fig. 7 shows the alpha power as a classification feature. The mean alpha power was obtained across all channels for this representation only. Each dot represents a healthy control participant or a PD patient (Blue: healthy controls; Orange: PD patients pre-exercise program; Green: PD patients post-exercise program). The amplitude alpha power was then used for training a simple classification.

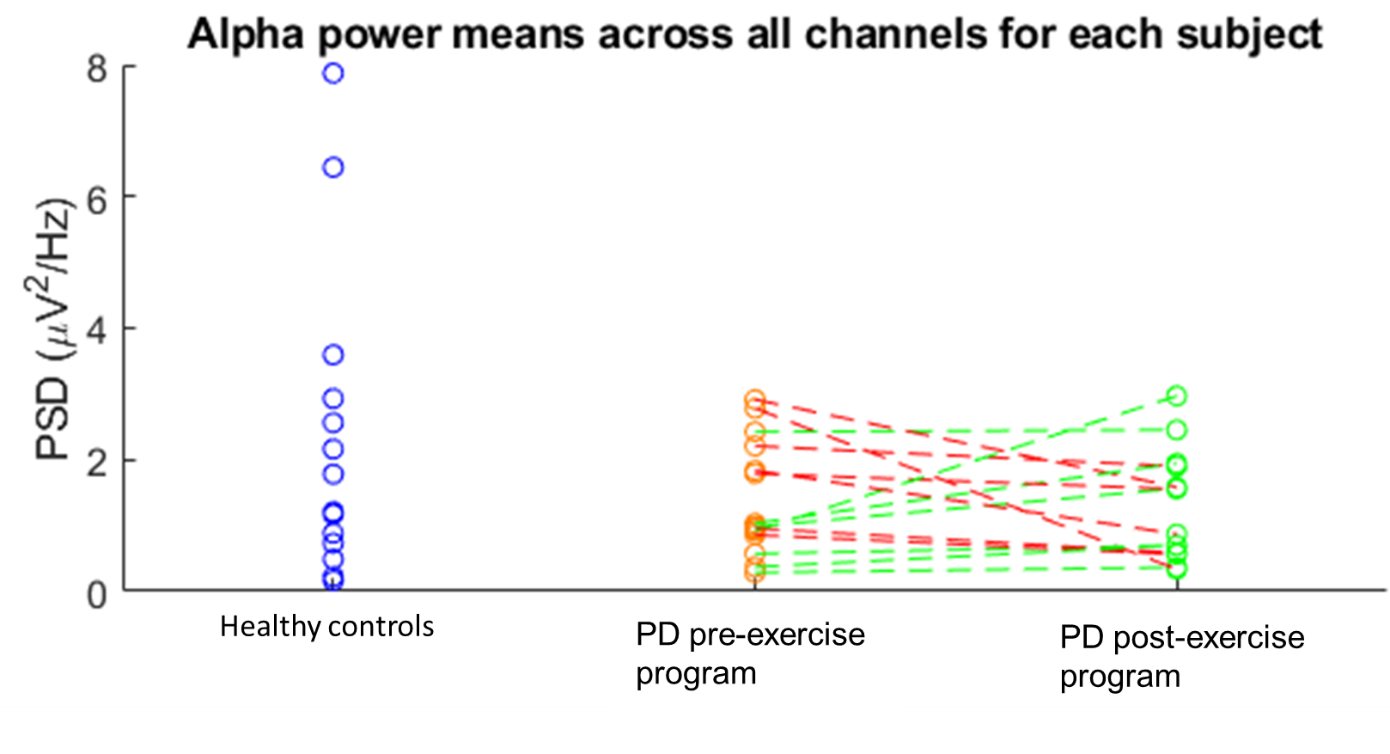

**Supp. Fig. 7.** Alpha power as a classification feature. The power spectrum density (PSD) for the alpha frequency band (8-14 Hz) was estimated from 2 min eyes closed RS-EEG data for each participant and patient in all channels. The mean alpha value was obtained across all channels for this representation only. Each dot represents a healthy control participant or a PD patient (Blue: healthy controls; Orange: PD patients pre-exercise program; Green: PD patients post-exercise program).

Supp. Fig. 8 shows the classification of healthy control participants and PD patients pre-exercise program using HsMM metrics as features (FP **top left**, DT mu **top right**, DT sigma **bottom right**); or the alpha power (**bottom left**) as features. For each set of features, a classifier was trained. Each dot represents a healthy control participant (blue) or a PD patient pre-exercise program (orange).

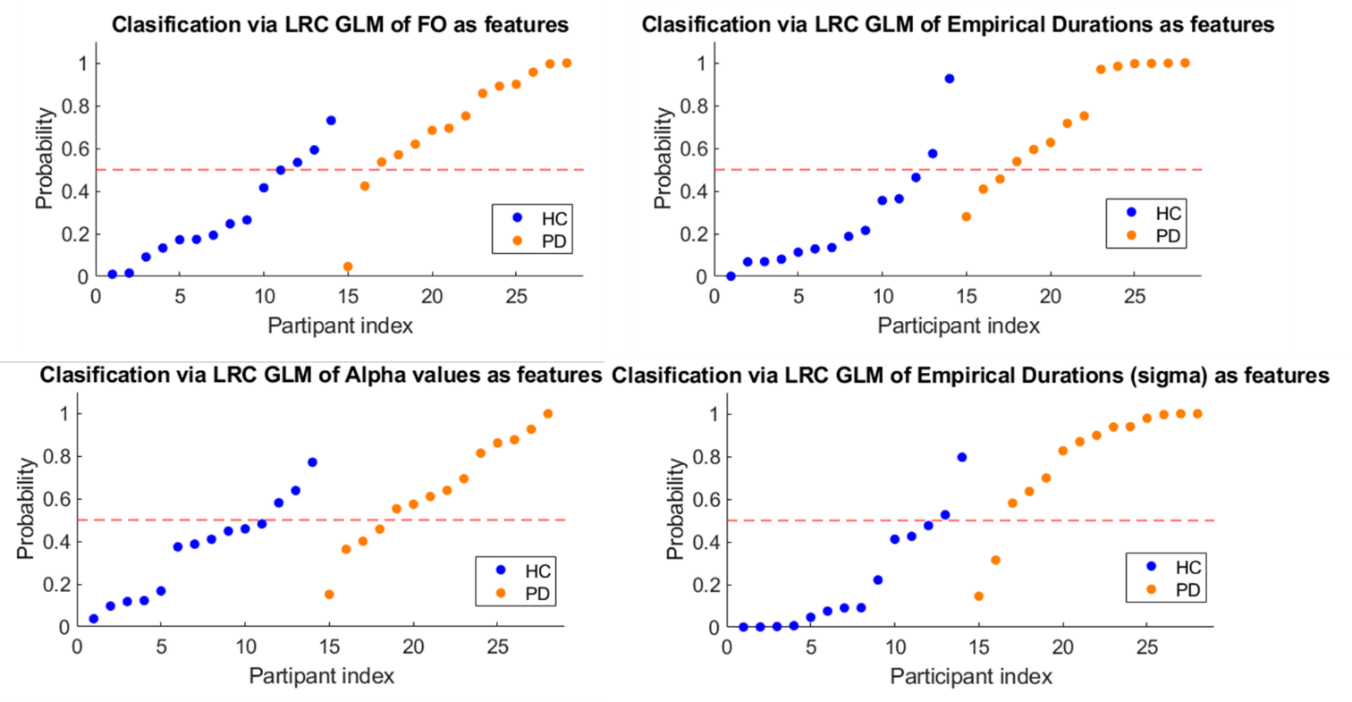

**Supp. Fig. 8.** Classification of healthy control participants and PD patients pre-exercise program using HSMM metrics (**Top left**, **top right**, **bottom right**) or alpha values (**bottom left**) as features. For each set of features, a classifier was trained using log-regression classification (LRC) based on General Linear Model (GLM). Each dot represents a healthy control participant (blue) or a PD patient pre-exercise program (orange).

In Supp. Fig. 9 it is shown each obtained ROC curve. This result shows the comparison of the performance of the classifiers based on HsMM dynamical features and the ones based on the RS-EEG alpha power. Three classifiers were trained using the dynamical features extracted from the HsMM model (FO blue line, DT *mu* red line, DT *sigma* yellow line). One classifier was trained using the alpha power as features (violet line). For each classifier the AUC was calculated (FO AUC: 0.8827; DT *mu* AUC: 0.9133; DT *sigma* AUC: 0.9286; Alpha AUC: 0.7908). Using a basic classifier for the feature sets we found that the best classification of participants and PD patients pre-exercise program is found for DT *sigma* as a feature, on the other extreme, the worse classification was obtained for alpha power as features. This tells us that the ability of HsMM dynamical features to characterize better the potential changes in activity between two independent groups or dependent groups.

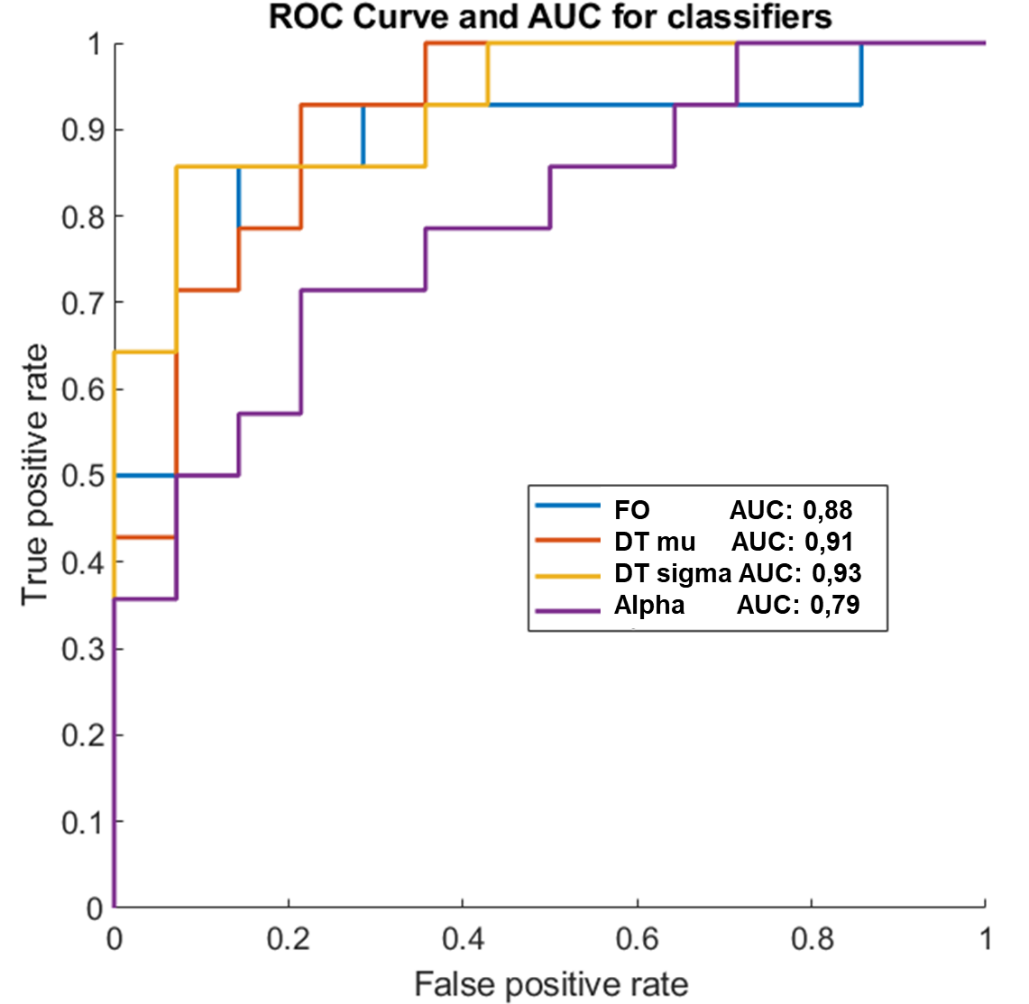

**Supp. Fig. 9.** Receiver operator characteristic (ROC) curves comparing the performance of classifiers based on dynamical features extracted from the HSMM to ones based on the RS-EEG alpha power values. Three classifiers were trained using the dynamical features extracted from the HSMM model (FO blue line, DT mean red line, DT variance, yellow line) and one classifier was trained using the alpha values as features (violet line). For each classifier the area under the curve (AUC) was assessed (FO AUC: **0.8827**; DT mean AUC: **0.9133**; DT variance AUC: **0.9286**; Alpha AUC: **0.7908**). Using a basic classifier for the feature sets we found that the best classification of participants and PD patients pre-training is found for DT variance as a feature, on the other extreme, the worse classification was obtained for alpha values band features. It is clear that the classifiers based on the dynamical features outperform classification based on the static spectral features.
